# The axillary lymphoid organ - an external, experimentally accessible immune organ in the zebrafish

**DOI:** 10.1101/2024.07.25.605139

**Authors:** Daniel Castranova, Madeleine I. Kenton, Aurora Kraus, Christopher W. Dell, Jong S. Park, Marina Venero Galanternik, Gilseung Park, Daniel N. Lumbantobing, Louis Dye, Miranda Marvel, James Iben, Kiyohito Taimatsu, Van Pham, Reegan J. Willms, Lucas Blevens, Tanner F. Robertson, Yiran Hou, Anna Huttenlocher, Edan Foley, Lynne R. Parenti, J. Kimble Frazer, Kedar Narayan, Brant M. Weinstein

**Author notes:** To whom correspondence should be addressed: Brant M. Weinstein, Section on Vertebrate Organogenesis, Building 6B, Room 4B413, 6 Center Drive, Bethesda, MD 20892. Current Address: University of Utah, Salt Lake City, UT, USA. These authors contributed equally (Madeleine I. Kenton, Aurora Kraus).

## Abstract

Lymph nodes and other secondary lymphoid organs play critical roles in immune surveillance and immune activation in mammals, but the deep internal locations of these organs make it challenging to image and study them in living animals. Here, we describe a previously uncharacterized external immune organ in the zebrafish ideally suited for studying immune cell dynamics *in vivo*, the axillary lymphoid organ (ALO). This small, translucent organ has an outer cortex teeming with immune cells, an inner medulla with a mesh-like network of fibroblastic reticular cells along which immune cells migrate, and a network of lymphatic vessels draining to a large adjacent lymph sac. Noninvasive high-resolution imaging of transgenically marked immune cells can be carried out in the lobes of living animals, and the ALO is readily accessible to external treatment. This newly discovered tissue provides a superb model for dynamic live imaging of immune cells and their interaction with pathogens and surrounding tissues, including blood and lymphatic vessels.

**Teaser:** A newly characterized external zebrafish lymphoid organ provides a powerful model for live imaging of immune cell dynamics

## Introduction

Vertebrates use tightly co-evolved adaptive and innate immune systems to combat pathogens. The innate immune system and its genetically encoded pathogen recognition receptors respond rapidly to invaders and their cytokine signals, helping tune the adaptive arm of the immune system to the type of pathogen present and influencing effector functions. Adaptive lymphocytes have highly specific somatically recombined receptors that, coupled to powerful effector functions, defend the host and impart memory for previously encountered pathogens. Pathogens are most frequently encountered at barrier tissues, where innate and adaptive immune cells work in concert to surveil the environment and protect the host.

Immune cells are born in primary lymphoid organs such as the bone marrow and thymus in humans and the head kidney and thymus in fishes, but secondary lymphoid organs (SLOs) are the locations where innate immune cells alert lymphocytes to pathogens and induce an effector response. In higher vertebrates such as humans the spleen, Peyer’s patches, and lymph nodes are sites where lymphocytes are activated by cognate antigens presented by antigen presenting cells (*1*). Germinal centers form in these SLOs when lymphocytes respond to antigen and proliferate, with B cells transcribing activation-induced cytidine deaminase (*aicda*) and undergoing affinity maturation with the help of T follicular helper cells to produce high avidity antibodies (*2*). Lymph nodes have a variety of characteristic features including a well-defined structure with an outer cortex and inner medulla, connection to draining lymphatic vascular networks, a mesh-like three-dimensional internal network of fibroblastic reticular cells (FRCs) facilitating immune cell trafficking, and specific sets of cytokines expressed by their resident cells that help direct lymphocyte homing and migration (*3–5*). Although fish lack organs that closely correspond to lymph nodes, they do have unique aggregations of adaptive lymphocytes where affinity maturation is thought to occur (*6*). In fishes, the head kidney (anterior kidney) and spleen are sites of adaptive immune activation, complete with the presence of antigen and *aicda* expression (*7, 8*). A number of recent studies have also identified mucosal associated lymphoid tissues (MALT) in bony fishes, including nasopharynx associated lymphoid tissue (O-NALT), diffuse gill associated tissues (GALT i.e. ILT, ALT and NELO) as well as a bursal tissue (*9–13*). The cellular mechanisms and mechanics of lymphocyte activation *in vivo* remain unclear in part because secondary lymphoid organs are extremely difficult to image intravitally in mammals.

The adult zebrafish *(Danio rerio)* is an emerging immune model that offers the ability to carry out high-resolution optical imaging of lymphocyte activation in living animals (*14*). In addition to a robust innate immune system, adult zebrafish have orthologous effector CD4 T cell subtypes, cytotoxic CD8 T cells, and B cells that produce IgM and mucosal IgT (*15*). They also have a lymphatic vascular system with many of the anatomical, molecular, and functional features of mammalian lymphatics (*16, 17*). While the known lymphoid organs of adult zebrafish are too deep within the organism to image, more externally located immune cell populations can be visualized in living adult animals, including a recently described tessellated lymphoid network (TLN) in the skin (*18*) although the anatomy of the TLN is not closely analogous to SLOs of mammals.

Here we report a previously uncharacterized external lobe located bilaterally just above the base of the pectoral fin. These small translucent external lobes contain a plexus of blood vessels and a network of lymphatic vessels draining to a large adjacent lymph sac. They have an outer cortex containing large numbers of immune cells, and an inner medulla with a fibroblastic reticular cell network and other features resembling lymph nodes and other secondary lymphoid organs, suggestive of a role in immune surveillance. Based on these and other features described below, we designated these lobes “Axillary Lymphoid Organs” (ALOs). ALOs can be imaged and are readily accessible to external treatment with antigens and pathogens in intact, living animals. They are also easily removed for *ex vivo* analysis, and they regenerate within a few weeks after removal. Together with a wide array of available zebrafish transgenic reporter lines marking numerous different immune, vascular, and other cell populations, this newly discovered organ provides a superb model for high resolution optical imaging of the interaction between immune cells, pathogens, and their surrounding tissues, including the vasculature.

## Results

### Anatomical characterization of the axillary lymphoid organ

Close examination of post-metamorphic zebrafish reveals a previously undescribed bilateral fleshy lobular structure immediately posterior to the operculum and just dorsal to the base of the pectoral fin, which we have denoted Axillary Lymphoid Organs (ALOs) based on the findings we report below. ALOs are small and translucent, most commonly lacking pigment cells, but they can be readily visualized by confocal microscopy after soaking adult fish in a fluorescent surface stain such as BODIPY^TM^ 630/650 (**Fig. 1A,B**). Although most appear as unilobular appendages (**Supp. Fig. 1A**), bilobed or multilobed ALOs are occasionally found in some animals (**Supp. Fig. 1B**), and rarely ALOs are found that do have pigment cells (**Supp. Fig. 1C**). ALOs emerge comparatively late in development, and are not found in embryonic or larval zebrafish. They first begin to make their appearance when fish reach around 10 mm in total length, at approximately 30 days post fertilization, with lobes increasing rapidly in size over the next few weeks (**Fig. 1D-G**). Like many other adult zebrafish organs and tissues, ALOs are able to completely regenerate after amputation, regrowing within about two weeks (**Fig. 1H-K, Supp. Movie 1**). Similar structures are found in other basal teleost fishes, including additional members of order Cypriniformes, which in addition to the zebrafish, includes Afro-Asian minnows (the families Danionidae and Xenocyprididae), loaches (the families Botiidae, Gastromyzontidae, Nemacheiliidae, and Vaillantellidae), and their relatives. A survey of fixed teleost specimens revealed a diversity of lobe morphologies, with some fish species possessing substantially larger or more elaborate structures than the ALOs of zebrafish (**Fig. 1L-O**, **Table 1**).

**Fig. 1.**
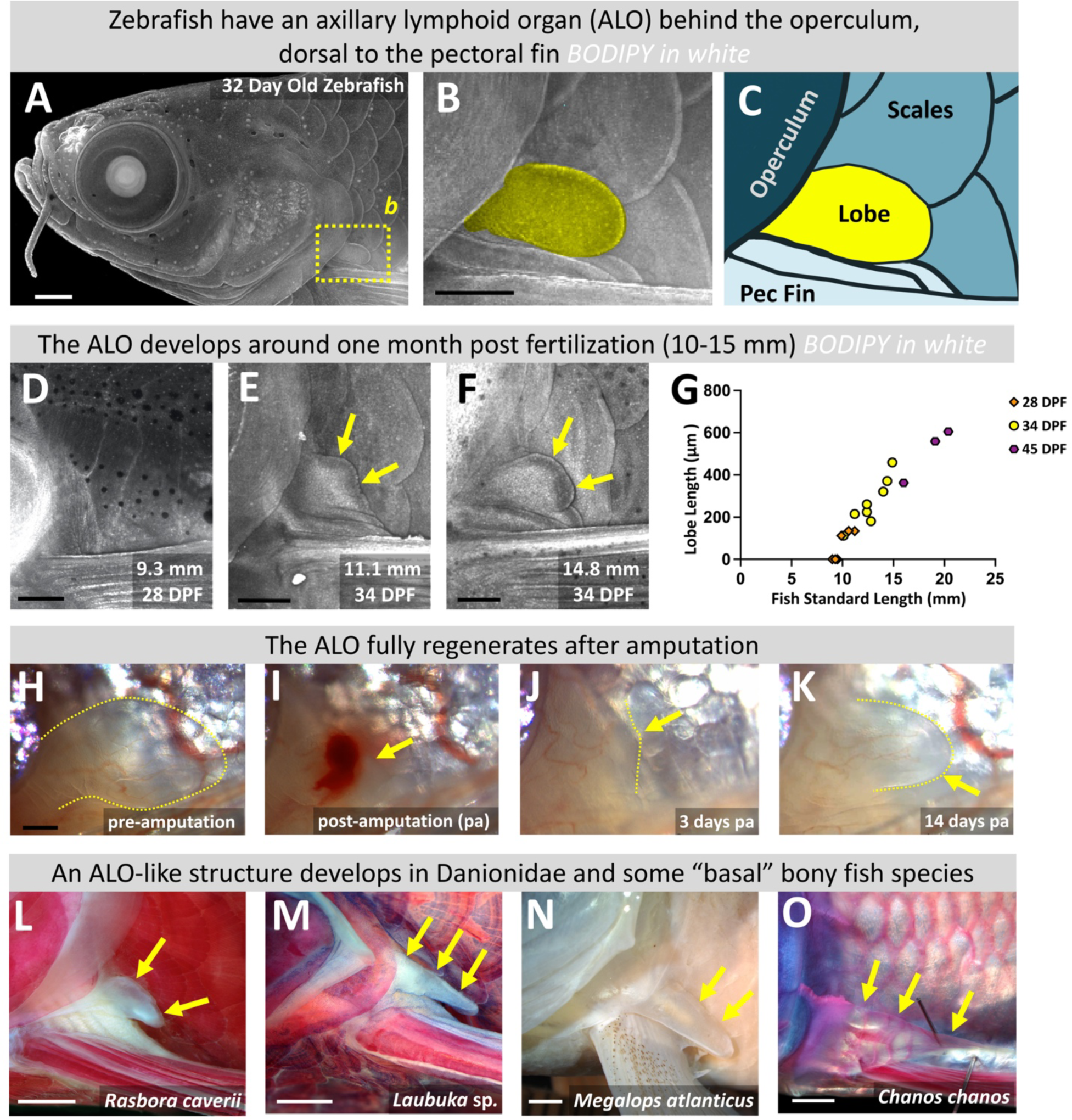
Axillary lymphoid organ (ALO) in the zebrafish. **A.** Maximum intensity projection confocal micrograph of the red fluorescent surface of a 32 day old zebrafish soaked in BODIPY 633. **B.** Magnified image of the boxed area in panel A, with the ALO pseudocolored yellow. **C.** Schematic diagram of the anatomical features shown in panel B. **D-F.** Maximum intensity projection confocal micrographs of 9.3 mm (28 dpf) (D), 11.1 mm (34 dpf) (E), and 14.8 mm (34 dpf) (F) juvenile zebrafish soaked in BODIPY 633, showing stages prior to ALO emergence, initial ALO budding, and further ALO expansion, respectively. **G.** Measurement of ALO length (µm) vs. fish standard length (mm). ALO’s emerge when fish reach approximately 9-10 mm in body length. **H-K.** Brightfield images of adult zebrafish ALO pre-amputation (H), immediately post-amputation (I), 3 days post-amputation (J), and 14 days post-amputation (K), with yellow dashed lines and arrows marking the border of the ALO. **L-O** Images of ALOs found on other fish species, with yellow arrows noting the locations of the ALOs. Scale bars = 500 µm (A,H), 250 µm (B), 200 µm (D,E,F), 1 mm (L,M), and 2mm (N,O)

To begin to examine the anatomical structure of the ALO, we collected transverse sections from fixed, paraffin-embedded whole adult zebrafish just caudal to the operculum, through the base of the pectoral fin (**Fig. 2A**). External, bilaterally located ALOs are readily observed just dorsal to the base of the pectoral fin in stained sections (**Fig. 2B**). Higher magnification images show that ALOs contain a clearly delineated and well separated cell-rich cortical region and a relatively cell-deficient matrix-rich medullary region (**Fig. 2C**). As noted above, most ALOs consist of a single lobe but fish with lobes containing multiple medullary compartments (**Supp. Fig. 1D,E**) or with multiple lobes (**Supp. Fig. 1F,G**) are also noted in histological sections.

**Fig 2.**
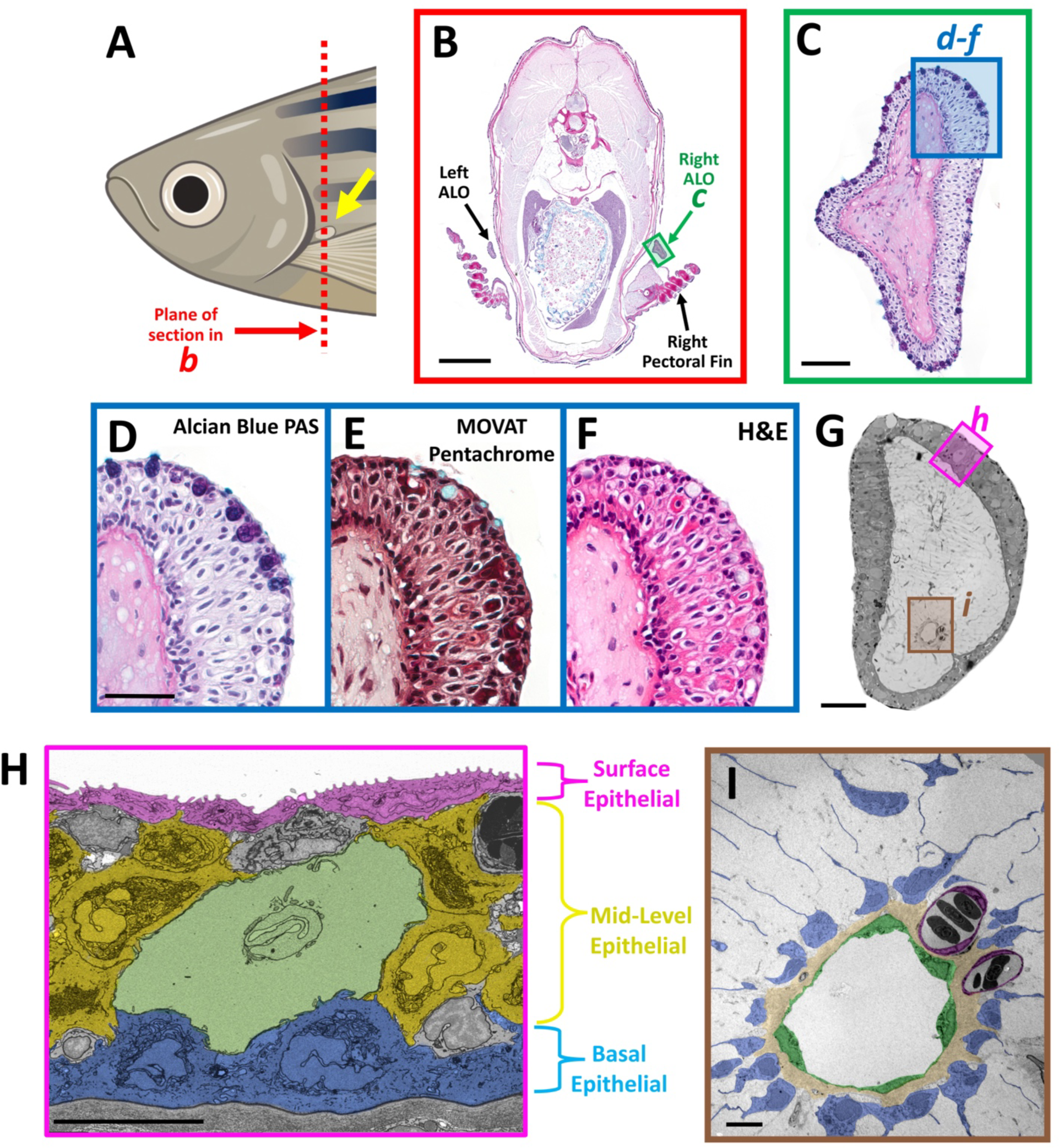
Histology and electron microscopy of the axillary lymphoid organ (ALO). Histological characterization of ALO morphology. **A.** Schematic diagram showing the plane of section in panel (B). **B.** Alcian Blue PAS-stained transverse paraffin section through the anterior trunk of an adult zebrafish. The green box indicates the region shown in panel C. **C.** Magnified image of the boxed region in panel B, showing the right ALO. The blue box indicates the region shown in panel D. **D-F.** Magnified images of serial transverse paraffin sections through the anterior trunk of an adult zebrafish stained with Alcian Blue PAS (D), Movat Pentachrome (E), and H&E (F). Panel D shows the boxed region in panel C. **G.** Transmission electron micrograph (TEM) of a transverse section through an adult zebrafish ALO. The boxed regions show areas with comparable (H) or actual (I) magnified TEM images in subsequent panels. **H.** Pseudocolored TEM showing the dermal cortex of the ALO with presumptive surface epithelial cells magenta, mid-level epithelial cells yellow, club cell green, and basal epithelial cells blue. **I.** Magnified TEM image of box I in panel G, showing the medulla of the ALO with lymphatic vessel pseudocolored green, blood vessels magenta, and fibroblastic reticular cells (FRCs) blue (**I**). Scale bars = 1 mm (B), 50 µm (C), 25 µm (D), 100 um (G), 10 µm (H,I).

Alcian Blue PAS, MOVAT pentachrome, and H&E staining of sectioned ALOs suggest that a variety of different cell types are present in the cortex in distinct surface, mid-level, and basal epithelial layers, including mucus-secreting goblet-like cells near the cortex surface identifiable by MOVAT pentachrome staining (*19*) (**Fig. 2C-F**). We also carried out transmission electron microscopy (TEM) and array tomography (AT) on ultrathin sections of ALOs for a more detailed morphological assessment of the structure and ultrastructure of the ALO and its constituent cells (**Fig. 2G-I, Supp. Figs. 1,3-5,7-9**). Electron microscopy of the ALO cortex confirms its division into surface, mid-level, and basal epithelial layers populated by cells displaying distinct morphologies (**Fig. 2H, Supp. Fig. 3**). The most superficial cells of the ALO cortex are flattened epithelial cells exposed to the outer environment with abundant numbers of “microridges,” actin-rich surface protrusions arranged in unique fingerprint-like patterns (**Supp. Fig. 1H-J**) that have been observed on surface skin cells of many different fish species (*20*). The mid-cortex contains mostly epithelial-like cells, although a few other cell types are also present, notably large cells resembling previously reported club cells (see below). The basal cortical layer contains mostly pyramidal or cuboidal epithelial cells with their apices pointing toward the outside of the ALO, and with their bases immediately apposed to a well-defined basement membrane separating cortical layers of the ALO from the deeper ALO medulla. The ALO medulla is composed largely of acellular matrix, but also contains unusual cells with thin, highly elongated, mostly radially arranged sheetlike reticular processes (**Fig. 2I**). Many but not all cell bodies of these medullary reticular cells are embedded in a collagenous matrix layer immediately adjacent to endothelial-lined lymphatic vessels that are distinguishable from blood vessels by their lack of red blood cells (**Fig. 2I**, and see below). In addition to TEM on individual ALO sections, we also carried out volume electron microscopy by array tomography to compile detailed 3-D ultrastructural visualization of ALOs and their cell populations (**Supp. Movie 2**, and see below).

### Single-cell analysis of the axillary lymphoid organ

To better understand the cellular composition of the ALO, we used the 10X Genomics Chromium platform to carry out single-cell RNAseq (scRNAseq) on a cell suspension prepared from 4 axillary lymphoid organs (2 from males, 2 from females) removed from adult zebrafish (**Fig. 3A**). An estimated total of 9,450 cells were sampled with 182,913,580 total sequence reads, with 93.9% of reads mapped to the genome and 76.4% of reads mapped to the zebrafish transcriptome. This represented 19,356 mean sequence reads per cell, 2,052 median UMI counts per cell, and 925 mean genes per cell (**Fig. 3B**). Unsupervised clustering using Seurat (*21*) identified 14 separate clusters that could all be definitively annotated as identifiable cell populations (**Fig. 3C**) based on their expression of characteristic cell-type specific genes, overall gene expression profiles, comparison to genes expressed in other single-cell data sets, notably Daniocell (*22*), and, importantly, spatial localization of cluster-specific transcripts to morphologically identifiable cells, all as discussed below.

**Fig 3.**
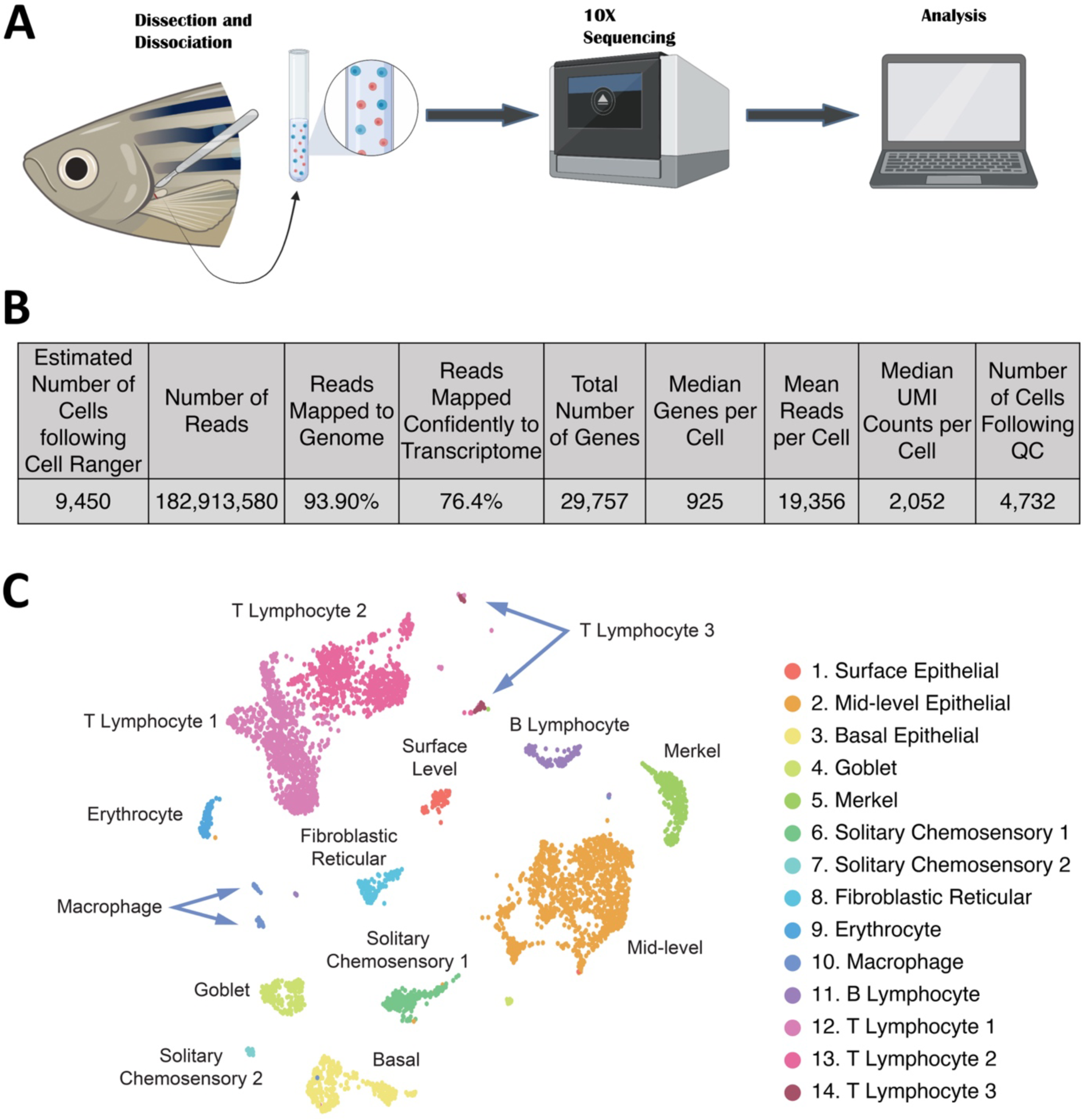
Single-cell RNA-seq of the ALO. **A.** Schematic diagram showing the workflow for ALO single cell RNA sequencing. **B.** Metrics for the ALO scRNA-seq procedure. **C.** Uniform Manifold Approximation and Projection (UMAP) plot of data from the ALO scRNA-seq procedure, with 14 clusters annotated by cell identity.

### Epithelial cells of the ALO cortex

Clusters 1-3 correspond to resident surface, mid-level, and basal epithelial cell populations in the ALO cortex (**Fig. 4A,B, Supp. Fig. 3A-C, Supp. Movie 2**), as confirmed by the methods noted above including *in situ* Hybridization Chain Reaction (HCR, (*23*)) using genes highly enriched in each of these cell clusters (**Supp Fig. 2**). Cluster 1 represents surface epithelium (**Fig. 4A,B**). This cluster specifically expresses c*laudin e (cldne)* and *krt1-19d* (**Supp Fig. 2C**), both of which are previously reported markers of surface epithelium in the zebrafish fin (*24*). HCR confirms that *cldne* expression is restricted to ALO surface epithelial cells (**Fig. 4C,D**). *dhrs13a.2, si:ch211-217k17.9,* and *ponzr2*, genes enriched in periderm in the Daniocell dataset (*22*), are also highly specific to this cluster (**Supp Fig. 2C**). These cells form a flattened epithelial monolayer on the surface of the ALO, with prominent characteristic apical microridges (*20*) and tight cell-cell junctions (**Fig. 4D, Supp. Figs. 1H-J, 3A-C, Supp. Movie 2**).

**Fig 4.**
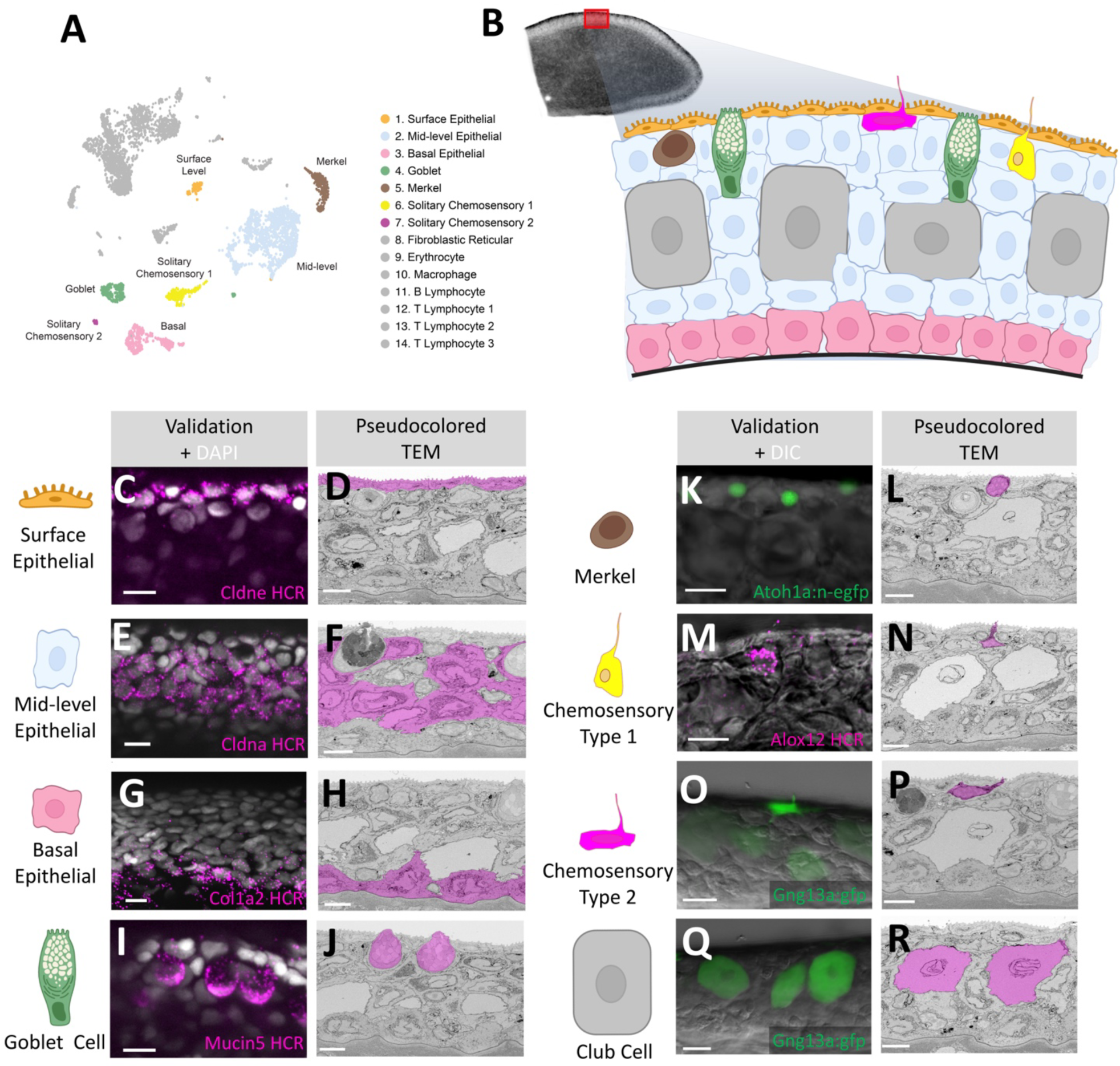
Resident cell types of the ALO dermal cortex. **A.** UMAP plot of ALO scRNA-seq data highlighting seven clusters that include resident cell types of the ALO cortex. **B**. Schematic diagram of the ALO cortex (comparable to the area marked by the red box in the ALO confocal image at upper left), with the cell types represented by each of the highlighted clusters in panel (A) shown using the same colors. **C,E,G,I,K,M,O,Q.** Confocal micrographs of the cortex of l hybridization chain reaction (HCR) stained (C,E,G,I,M) or transgene-expressing (K,O,Q) ALOs isolated from adult zebrafish. **D,F,H,J,L,N,P,R.** Pseudocolored 2D sections from an array tomography stack of the ALO cortex with the same cell types shown in the adjacent confocal image panels highlighted in pink. The confocal images and electron micrographs show surface epithelial cells (**C,D**), mid-level epithelial cells (**E,F**), basal epithelial cells (**G, H**), goblet cells (**I, J**), Merkel cells (**K, L**), chemosensory type 1 cells (**M, N**), chemosensory type 2 cells (**O, P**), and club cells (**Q, R**). Scale bars = 10 µm (C, E, G, I, K, M, O, Q) or 5 µm (D, F, H, J, L, N, P, R).

Cluster 2 corresponds to mid-level or intermediate cortical epithelial cells (**Fig. 4A,B, Supp. Fig. 3A,D-F, Supp. Movie 2**). The previously characterized intermediate epidermal marker *claudin a (cldna)* (*24*) is highly enriched in this cluster (**Supp. Fig. 2C**), and HCR confirms *cldna* expression is localized to ALO mid-level epithelial cells (**Fig. 4E,F**). Other genes highly enriched in this cluster include *epidermal growth factor receptor a (egfra)*, a gene involved in positive regulation of epithelial cell proliferation and inflammatory/immune reactions in mammalian skin (*25*), and *foxq1a*, a gene found in periderm in the developing zebrafish (*22*) (**Supp Fig. 2C**). Mid-level epithelial cells have a varied appearance and morphology in TEM images (**Fig. 4F, Supp. Movie 2**), although most have abundant cell-cell adherens junctions (**Supp. Fig. 3D,E**) and intracellular fibrillar networks (**Supp. Fig. 3D,F**).

Cluster 3 contains basal epithelial cells of the ALO cortex (**Fig. 4A,B, Supp. Fig. 3A,G-I, Supp. Movie 2**). Previously identified zebrafish tail fin basal epidermal markers *col1a2* and *krtt1c19e* (*24*) are highly enriched in this cluster (**Supp Fig. 2C**), and HCR shows *col1a2* is restricted to the deepest epithelial layer of the ALO cortex, adjacent to the basal lamina (**Fig. 4G,H**). Basal epithelial cells also specifically express epidermal-enriched *itgb4* and *epgn* genes (*22*), further confirming their epithelial identity (**Supp Fig. 2C**). *Epgn (epigen* or *epithelial mitogen*) is predicted to enable epidermal growth factor receptor (EGFR) binding activity and growth factor activity, and this cluster is enriched for genes involved in Rho GTPase signaling. Many basal epithelial cells have a roughly pyramidal morphology (**Fig. 4H, Supp. Movie 2**), with a centrally located nucleus with abundant adjacent golgi localized to the externally-facing side of the nucleus. The basal epithelial cell layer is immediately adjacent to a well-defined basal lamina (**Supp. Fig. 3G,H**, red arrows) and underlying thick extracellular matrix layers that separate the ALO cortex from the ALO medulla, and together they form a monolayer with abundant tight cell-cell junctions (**Supp. Fig. 3G,I**, yellow arrow).

### Additional resident cell types of the ALO cortex

In addition to epithelial cells, several other resident cell types are also present in the ALO cortex (**Fig. 4A,B, Supp. Figs. 4-8**). Cluster 4 represents mucus-producing goblet cells (**Fig. 4I,J, Supp. Fig. 4**), readily identified by their specific expression of the *muc5.1* and *muc5.2* mucin genes (*26*) (**Supp. Figs. 2C, 4**). Mucins are expressed in intestinal goblet cells (*27*) and in analogous epidermal mucous-producing cells of other organs and tissues such as the respiratory and reproductive tracts, and in fish skin (*26*), where they play an important role in protection from infection. HCR for *muc5.1* confirms expression of this gene localizes to morphologically distinctive goblet-like cells that display characteristic ultrastructural features of this cell type including internal contents spilling out from the ALO surface (**Fig. 4I,J, Supp. Fig. 4A-E, Supp. Movies 2,3**). Cluster 4 also strongly and specifically expresses *p2rx1*, *fer1l6*, and *si:dkey-65b12.6* (**Supp. Fig. 2C**), each of which is highly enriched in epidermal mucous secreting cells in the Daniocell scRNAseq data set (*22*). The *agr2* gene is most highly expressed in cluster 4, although it is also weakly expressed in clusters 6 and 14 (see below). *Agr2* is a protein disulfide isomerase found in mammalian intestinal goblet cells important for proper processing of gel-forming mucins in humans (*28*). Cluster 4 is also the cluster most strongly expressing interleukin 13 receptor *il13ra1*, although again it is expressed at lower levels in clusters 6, 7, and 11 (see below). IL-13 is a cytokine known to stimulate goblet cell differentiation and goblet cell mucus production (*29*). In addition to their passive role in immune defense via mucus production, goblet cells may also play more active roles in immunity, including taking up and presenting antigens to underlying antigen-presenting cells that induce adaptive immune responses (*30*). Interestingly, zebrafish ALO goblet cells also possess a robust capacity to take up exogenous substances (**Supp Fig. 4F-H**).

Cluster 5 consists of Merkel cells (**Fig. 4A,B, Supp. Fig. 5**), as confirmed by their previously described highly specific expression of an *atoh1a:nls-Eos* transgene (*31*), and by their characteristic location and morphology (*31, 32*) (**Fig. 4K,L**). Merkel cells are mechanosensory cells sensitive to gentle touch stimulation known to express neuroendocrine markers such as *chromogranin-a (chga)*, which like *atoh1a* is also highly expressed in and highly specific for ALO cluster 4 (**Supp Fig. 2C**). Cluster 4 cells also express neural- and sensory-associated genes such as *kcnd2, srrm4,* and *penka* (**Supp Fig. 2C**). Like their mammalian counterparts (*33*), zebrafish Merkel cells have a number of characteristic morphological and ultrastructural features including their location just under the surface epithelium, small size and relatively round cell shape, and ventral cellular projection (**Supp Fig. 5, Supp. Movie 2**).

Clusters 6 and 7 correspond to two classes of solitary chemosensory cells (SCCs) (**Fig. 4A,B, Supp. Figs. 6 and 7, Supp. Movie 3**). Cluster 6 Type 1 “tuft-like” SCCs (SCC1) show highly specific expression of known tuft cell markers such as *avil* and leukotrienes *alox5a and alox12* (**Supp Fig. 2C**). ALO HCR shows *alox12* is expressed in cells with cell bodies located just under the surface epithelium that have sensory-like apices protruding out through the epithelium (**Fig. 4M,N, Supp Fig. 6C,D**). Tuft cells sense external stimuli at mucosal barriers, responding by secreting effector molecules such as leukotrienes capable of evoking neural, immune, and other responses (*34*). Cluster 7 Type 2 “taste-like” SCCs (SCC2) show high expression of *gng13a* and *id2b,* which are highly expressed in taste cells in the Daniocell data set (*22*) (**Supp Fig. 2C**). Cluster 7 SCC2 cells are specifically marked by robust expression of a *gng13a:egfp* transgene (**Fig. 4O,P, Supp Fig. 6A,B,E-G**). The two SCC cell types also share expression of several characteristic genes such as *sox8b* and *plcg2* (**Supp Fig. 2C**). SCC cells with apical extensions penetrating through the surface epithelium were readily identified by DIC imaging (**Supp Fig. 6D,G**). SCC-like cells were also observed in TEM and array tomography images of the ALO (**Supp Fig. 7, Supp. Movie 3**), although ultrastructural subtypes corresponding to the *alox12* HCR-positive (SCC1) and *gng13a* HCR- or *gng13a:egfp* transgene-positive (SCC2) cell clusters could not be readily distinguished. Interestingly, TEM array tomography showed that SCC-like cell invariably had two separate apical projections protruding through the surface epithelium, generally positioned on opposite ends of the cell (**Supp Fig. 7, Supp. Movie 3**).

In addition to the seven cortical cell types corresponding to identified scRNAseq clusters noted above, histological and ultrastructural imaging of the ALO also revealed abundant numbers of large, mostly ovoid cells located within the mid-level epithelium (**Fig. 4B,R, Supp Fig. 8A, Supp. Movie 2**). These large cells also have a homogeneous-appearing cytoplasm and centrally located, sometimes binucleate or more complex-shaped nuclei (**Fig. 4R, Supp. Fig. 8A-C, Supp. Movie 2**), all features indicative of club cells, an unusual cell type described in mammalian lungs (*35*) and in fish skin (*36*), including in zebrafish (*37–39*). In fish, club cells are reservoirs for a “fright substance” (Schreckstoff) released when fish are injured that elicits fear responses in nearby conspecific animals (*38*), and they have also been implicated in innate or acquired immunity (*37, 39*). HCR staining with genes specific for each of the clusters identified in our single cell analysis (**Fig. 3C**) failed to mark the club cell population, suggesting that these large cells were not captured in our single cell analysis. Interestingly, confocal imaging shows that these cells are very weakly positive for the same *gng13a:egfp* transgene that strongly marks SCC2 cells, allowing them to be visualized by confocal imaging (**Fig. 4O,Q, Supp. Fig. 8D-F**).

### Vasculature of the ALO medulla

As noted above, the highly cellular, densely packed ALO cortex surrounds a medulla composed of a largely acellular matrix with a sparsely interspersed network of highly elongated cells (**Fig. 2**). Although vasculature is absent from the ALO cortex, the medulla contains networks of blood and lymphatic vessels readily visualized using *Tg(kdrl:mcherry)^y206^*and *Tg(mrc1a:eGFP*)*^y251^* transgenic reporter lines (*40, 41*) labeling blood (magenta) and lymphatic (green) endothelium, respectively (**Fig. 5A-C**). Circulating nucleated zebrafish red blood cells marked by intravascular injection of Hoechst nuclear stain into the caudal axial vasculature (blue) circulate robustly though the blood vessels (magenta) but not through the lymphatic vessels (green), confirming their identification (**Fig. 5B-D, Supp. Movie 4**). The identification of lymphatic vessels is further confirmed by injection of Qtracker^TM^ 705 vascular labels (quantum dots) into the ALO medulla (**Fig. 5E**). The quantum dots (magenta) are specifically taken up by the lymphatic vessels and drained efficiently into a large lymph sac (arrow in panel G) located nearby but deeper in the body of the fish (**Fig. 5F,G**). Blood and lymphatic vessels can both be visualized ultrastructurally in TEM images of the ALO medulla (**Fig. 2G,I**) with characteristic tight junctions (**Fig. 5H-I**). Vessels are enveloped by a matrix-rich layer and lymphatic vessels in particular are surrounded by numerous cell bodies of unusual medullary reticular cells (**Fig. 2G,I**), described below.

**Fig. 5.**
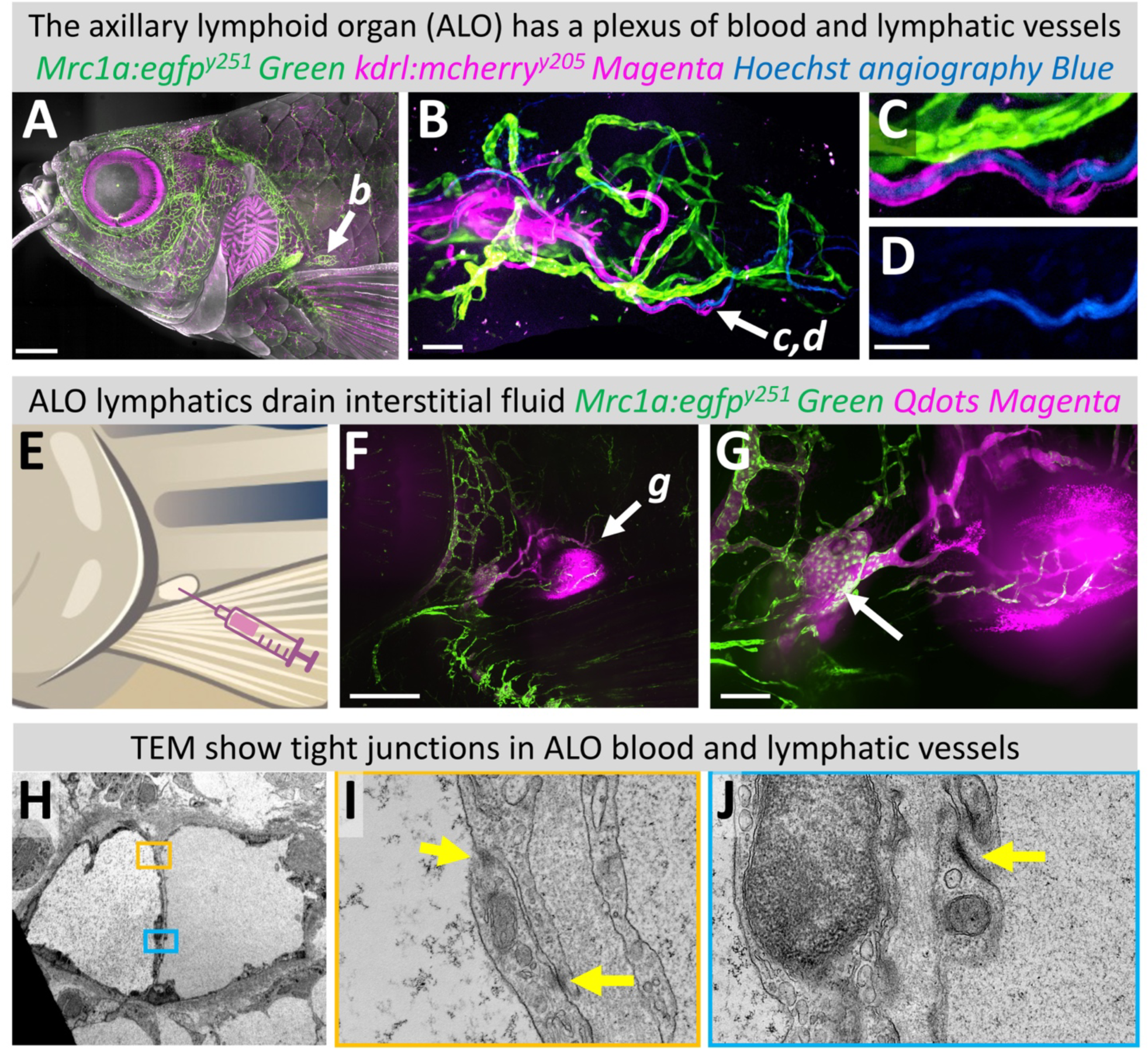
The ALO contains a network of blood and lymphatic vessels. **A-D.** Confocal micrographs of an adult *Tg(mrc1a:egfp)^y251^, Tg(kdrl:mcherry)^y205^* double transgenic zebrafish with mrc1a-positive lymphatics in green and kdrl-positive blood vessels in magenta. This animal was also injected intravascularly with Hoechst 33342 dye, marking blood cell nuclei with blue fluorescence. Images include (A), an overview image of the head, with the position of the ALO noted, (B), higher magnification image of the area in panel A noted by an arrow, showing a network of blood (magenta) and lymphatic (green) vessels in the ALO, with blood circulation (blue) in blood vessels, (C,D), higher magnification images of the area noted by an arrow in panel B, showing that blood vessels (magenta) but not lymphatics (green) contain circulating RBCs (blue). **E.** Schematic diagram of procedure for intralobular injection of quantum dots. **F,G.** Confocal micrographs of an adult *Tg(mrc1a:egfp)^y251^* zebrafish ALO after intra-organ injection of 705 nm quantum dots (Qdots), with mrc1a-positive lymphatics in green and Qdots in magenta. Panel F shows an overview of the ALO vicinity, noting the higher-magnification area shown in panel G. The injected Qdots drain via lymphatics into an adjacent deeper large lymph sac, noted with an arrow in panel G. **H-J.** Transmission electron micrographs of ALO vessels, showing overview of adjacent vessels (H) and higher magnification images of vessel cell-cell junctions (I,J). Scale bars = 1 mm (A), 500 µm (F), 100 µm (G), 50 µm (B), 25 µm (D).

### Fibroblastic reticular cells of the ALO medulla

In addition to blood and lymphatic vessels, the medullary core of ALOs contains an elaborate, mesh-like network of cells linked together by long, thin tortuous sheetlike cellular projections that resemble FRC networks found in mammalian lymph nodes and other lymphoid organs, where they help to organize and direct immune cell migration and activity (*4, 5*) (**Figs. 2G,I, 6, Supp. Fig. 9, Supp. Movie 5**). ALO scRNAseq reveals a single distinct cluster (cluster 8) that strongly expresses *platelet-derived growth factor receptor beta (pdgfrb)* and *vimentin (vim),* both highly enriched in mammalian FRCs (*4*) (**Fig. 6A,B**), and confocal imaging of ALOs in a *TgBAC(pdgfrb:egfp)^ncv22^* transgenic reporter line (*42*) shows that FRCs are strongly EGFP-positive (**Fig. 6C-E, Supp. Fig. 9A,B, Supp. Movie 5**). Cluster 8 also strongly expresses *spock3,* a gene recently associated with dendritic cells in the developing zebrafish thymus (*43*), and *spock3* HCR specifically marks medullary FRCs in zebrafish ALOs, further validating the identity of these cells (**Fig 6B,F,G**). In mammalian secondary lymphoid organs, FRCs express chemokine ligands that provide guidance cues to direct and organize lymphocyte migration (*4*). Zebrafish ALO FRC cluster 8 similarly strongly expresses several different chemokines including *cxcl12a* and *ccl25b* (**Fig. 6B**) which are likely signaling to different immune cell populations present in the ALO expressing cognate chemokine receptors (see below). Live DIC imaging of the ALO medulla shows macrophages and other presumptive immune cells migrating actively along FRCs (**Fig. 6H, Supp. Movie 6**), and TEM images show close association between FRCs (blue) and presumptive immune cells (purple) (**Fig. 6I,J**). Interestingly, most FRC cell bodies are closely juxtaposed to lymphatic vessels and the matrix surrounding these vessels (**Fig. 6K, Supp. Fig.9C-E, Supp. Movie 6**), suggesting there may also be communication and cellular interaction between FRCs and lymphatic endothelial cells.

**Fig. 6.**
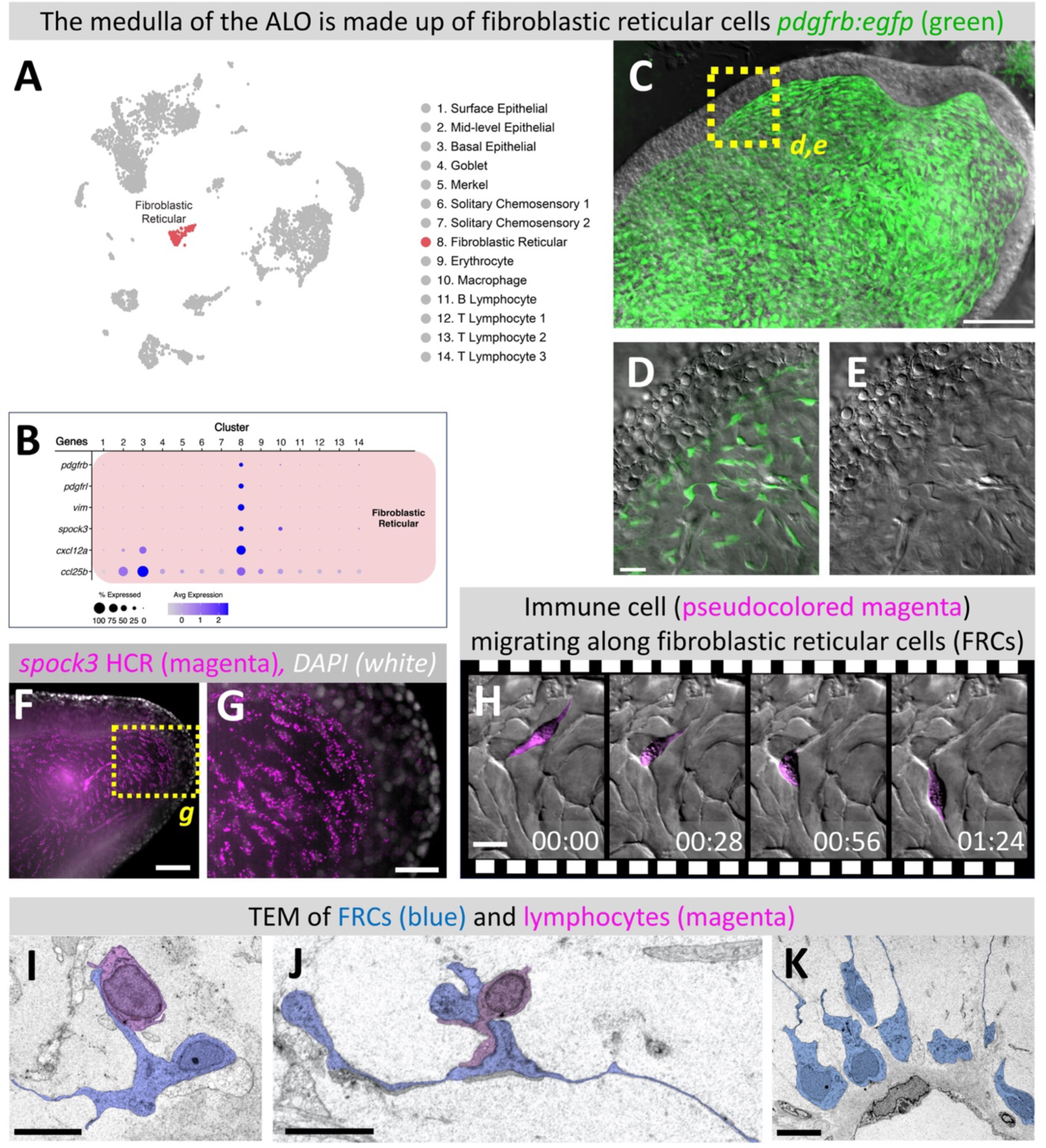
The ALO medulla is made up of fibroblastic reticular cells. **A.** UMAP plot of ALO scRNA-seq data highlighting the medullary fibroblastic reticular cell (FRC) cluster. **B.** Dot plot showing the relative gene expression of genes used to identify and characterize the FRC cluster. **C-E**. Confocal (C,D) and DIC (E) micrographs of an ALO excised from an adult *Tg(pdgfrb:EGFP)^ncv22^* transgenic zebrafish with green fluorescent FRCs. Panel C shows an ALO overview image, panels D and E show higher magnification images of the yellow boxed area in panel C. **F-G.** Confocal micrographs of the ALO of an adult zebrafish subjected to hybridization chain reaction (HCR) for *spock3* (magenta), with DAPI counterstain shown in white. Panel G shows a higher magnification image of the boxed area in panel F. **H.** Successive images from a timelapse DIC videomicrograph of an immune cell (pseudocolored magenta) migrating along the FRC network. Images are selected frames from **Supplemental Movie 6**. **I-K**. Transmission electron micrographs of the ALO medulla. Panels J and K show immune cells (pseudocolored magenta) in close apposition to FRCs (pseudocolored blue). Panel K shows FRC cell bodies adjacent to and embedded in the matrix surrounding a lymphatic vessel. Scale bars = 100 µm (C), 50 µm (F), 20 µm (D, G), 10 µm (H), 5 µm (I, J, K).

### The zebrafish axillary lymphoid organ contains large numbers of immune cells

In addition to epithelial, sensory, and stromal cells, scRNAseq of the ALO reveals large numbers of hematopoietic cells in clusters 9-14, including erythrocytes, macrophages, B cells, and several distinct classes of T cells **Fig. 7A,B**). Cluster 9 cells are easily identified as erythrocytes by their specific expression of hemoglobins *hbaa1* and *hbba1,* as well as other genes (**Fig. 7B**; fish erythrocytes are nucleated and transcriptionally active).

**Fig. 7.**
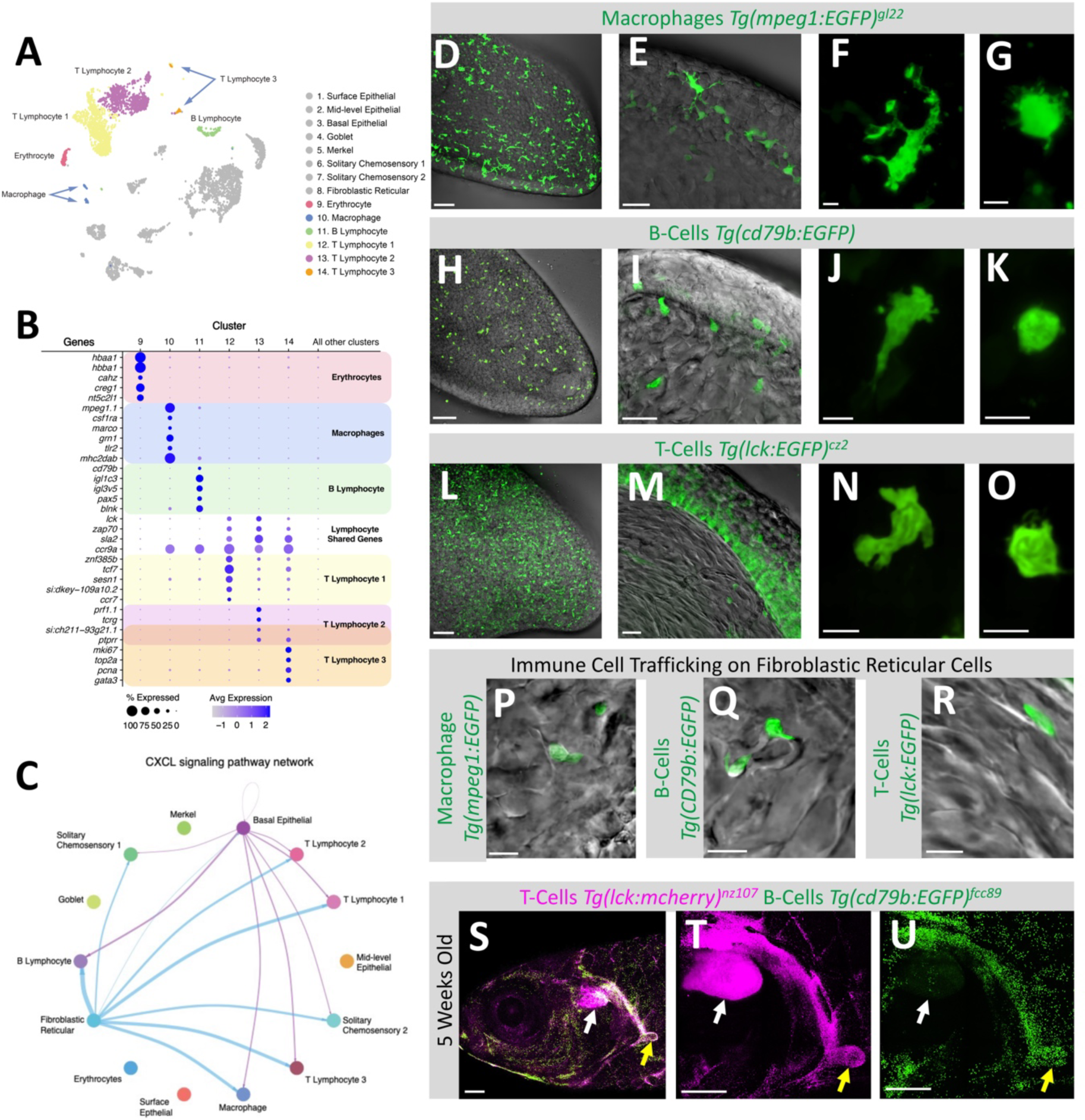
The ALO is an immune cell hub. **A.** UMAP plot of ALO scRNA-seq data highlighting blood and immune cell clusters. **B.** Dot plot showing the relative expression of genes used to identify and characterize blood and immune cell clusters. **C.** CellChat plot of likely chemokine signaling in the ALO based on expression of chemokine ligands and receptors in different ALO scRNAseq clusters. Basal epithelial and fibroblastic reticular cells both appear to be hubs for chemokine signaling to immune cells. **D-O.** Confocal + DIC (D,E,H,I,L,M) or confocal only (F,G,J,K,N,O) micrographs of ALOs from adult immune cell transgenic reporter zebrafish. Images include ALO overviews (D,H,L), side views of the dermal cortex and medulla (E,I,M), and higher magnification images of individual cells (F,G,J,K,N,O). Images show *Tg(mpeg1:EGFP)^gl22^* positive macrophages (D-G), *Tg(cd79b:EGFP)^fcc89^* positive B-cells (H-K), and *Tg(lck:EGFP)^cz2^* positive T-cells (L-O). **P-R**. Single-plane confocal + DIC images of *Tg(mpeg1:EGFP)^gl22^*positive macrophages (P), *Tg(cd79b:EGFP)^fcc89^* positive B cells (Q), and *Tg(lck:EGFP)^cz2^* T-cells (R) migrating on fibroblastic reticular cells in adult ALO medullae (see **Supp. Movies 6-9**). **P-R.** Confocal micrographs of a 5 week-old *Tg(cd79b:EGFP)^fcc89^*, *Tg(lck:mcherry)^nz107^* double transgenic zebrafish with T-cells in magenta and B-cells in green, showing large numbers of cells migrating between the thymus (white arrow) and the ALO (yellow arrow). Images show an overview of the head (S) and higher magnification images of T cells (T) connecting the thymus and ALO and B cells (U) starting above the thymus and moving down to the ALO (see **Supp. Fig. 10** for whole-fish overview images). Scale Bars = 50 µm (D, H, L), 20 µm (E, I, M), 5µm (F, G, J, K, N, O), 10 µm (P, Q, R), and 500 µm (S, T, U).

Cluster 10 specifically expresses many well-known diagnostic markers of macrophages, including *macrophage expressed gene 1.1 (mpeg1.1), colony-stimulating factor 1 receptor (csf1ra), macrophage receptor with collagenous structure (marco),* and *granulin 1 (grn1)* (**Fig. 7A,B**). The toll-like receptor 2 (tlr2) gene is also specifically expressed in the macrophage cluster, in alignment with the role of macrophages as pathogen detectors (**Fig. 7B**). Cluster 10 macrophages also strongly express *mhc2dab*, reflecting their function as antigen presenting cells (**Fig. 7B**). Confocal imaging of a *Tg(mpeg1.1:egfp)^gl22^*transgenic line (*44*) shows that ALO macrophages are found primarily in the cortex (**Fig. 7D-G**), although an occasional macrophage can be seen in the medulla migrating along FRCs (**Fig. 7P, Supp. Movie 6**). Live timelapse imaging of *Tg(mpeg1.1egfp)^gl22^* transgenics reveals two distinct phenotypes - highly motile cells that can be observed moving between the cortex and medulla, and less motile cells in the cortical layer of the ALO with many cellular extensions protruding in all directions (**Fig. 7F,G, Supp. Movie 7**). CellChat (*45*) mapping of CXCL signaling based on our scRNAseq data suggests that FRCs and basal epidermal cells both serve as major hubs for signaling to macrophages (**Fig. 7C**), and lymphocytes (see below).

Clusters 11-14 include several different lymphocyte cell populations (**Fig. 7A,B**). Cluster 11 is a small cluster of B lymphocytes, expressing various immunoglobulin components *(cd79b, igl1c3, igl3v5)* as well as *pax5,* known to mark commitment to B cell fate in mammals, and *blnk,* known to be a part of B cell activation signaling (**Fig. 7B**) (*46, 47*). Confocal imaging with a *Tg(cd79b:egfp)^fcc89^* transgenic B cell reporter line (*48*) shows B lymphocytes scattered throughout the cortex and medulla at homeostasis (**Fig. 7H,I**). Live time lapse imaging of *cd79b-*positive B lymphocytes reveals multiple distinct cellular behaviors (**Supp. Movie 8**). Some B lymphocytes migrate rapidly, sending numerous cellular protrusions in all directions (**Fig. 7J, Supp. Movie 8**) (*49*), while other B lymphocytes remain more spherical and stationary, although they can still be observed continuously extending smaller filopodia (**Fig. 7K, Supp. Movie 8**). B lymphocytes are also readily observed migrating actively on the FRC network (**Fig. 7Q, Supp. Movie 8**).

Clusters 12-14 represent different subsets of T lymphocytes, as confirmed by their common expression of diagnostic markers such as T cell receptor complex genes *lck* and *zap70* (**Fig. 7A,B**). Cluster 12 (T Lymphocyte 1) expresses transcription factor *tcf7,* known to be present in both naïve and memory T cells (*50*), while cluster 13 (T Lymphocyte 2) specifically expresses genes associated with cytotoxic lymphocytes, including *perforin1.1* (*prf1.1*) and *t cell receptor gamma*, *(tcrg)* (**Fig. 7B**). The final small cluster of cells, cluster 14 (T Lymphocyte 3) appears to be proliferating lymphocytes based on the expression of many genes associated with cell cycle activation (*mki67, topa, pcna*). A small subset of all three T lymphocyte clusters also expresses *ccr7*, a gene associated with migratory T cells (*51*). Confocal imaging of the ALO using T lymphocyte-specific *Tg(lck:egfp)^cz2^*reporter fish (*52*) reveals strikingly enriched localization of these cells near the basal cortex of the ALO (**Fig.7L-M, Supp. Movie 9**), consistent with CellChat identification of chemokine signaling between ALO basal epithelial cells and lymphocyte cell populations (**Fig. 7C**). Live time lapse imaging of *Tg(lck:egfp)^cz2^*ALOs shows that *lck*-positive lymphocytes are highly active, rapidly moving cells (**Supp. Movie 9**). Many cells have elongated cell shapes consistent with streaming T lymphocytes, but even more spherical T cells display numerous active filopodial protrusions (**Fig. 7N,O**).

As noted above, immune cells including macrophages, B cells, and T cells closely associate with and actively migrate along the sheetlike FRC network in the central ALO medulla (**Figs. 6H-J, 7P-R, Supp. Movies 6-9**). CellChat analysis of our RNAseq data indicates that FRCs serve as a chemokine signaling hub for all these immune cell types (**Fig. 7C**). FRC cell bodies are most highly concentrated along lymphatic vessels (**Figs. 2I, 6K, Supp. Movie 5**), which are themselves conduits for immune cell trafficking, suggesting FRCs serve as a pathway for immune cell trafficking between ALO lymphatics and the ALO cortex. Interestingly, imaging of a *Tg(lck:mcherry)^nz107^, Tg(cd79b:egfp)^fcc89^* double transgenic zebrafish (T cells red, B cells green) at 5 weeks, when many naïve T cells are leaving the thymus, also reveals a dense path of T cells directly connecting the thymus (white arrow) and the ALO (yellow arrow) (**Fig. 7S,T, Supp. Fig. 10A,B**), consistent with a role for the ALO as an immune surveillance hub. This pathway is also used by large numbers of B cells (**Fig. 7S,U, Supp. Fig. 10A,C**).

### T cell leukemia infiltrates the axillary lymphoid organ

In further support of its potential role as an immune nexus, the axillary lymphoid organ becomes highly infiltrated with leukemic cells in zebrafish leukemia models (**Fig. 8, Supp. Movie 10**). Previous studies using an established zebrafish model of T cell acute lymphoblastic leukemia (T-ALL) have revealed accumulation of lymphocytes in known secondary lymphoid organs such as the head kidney marrow and gill-associated lymphoid tissue (*53*). In the HLK^dz102^ model of T-ALL, axillary lymphoid organs frequently become preferentially and massively infiltrated with GFP-positive leukemic T cells (**Fig. 8A,B**). Leukemic T cells accumulate in large numbers both in the cortex, where substantial numbers of T cells are present in wild type animals, and in the medulla, where there are usually fewer actively migrating T cells (**Fig. 8C-F**). Eventually, infiltrating leukemic cells completely fill most of the ALO, displacing many other cells in the cortex (**Fig. 8G-J, Supp. Movie 10**).

**Fig. 8.**
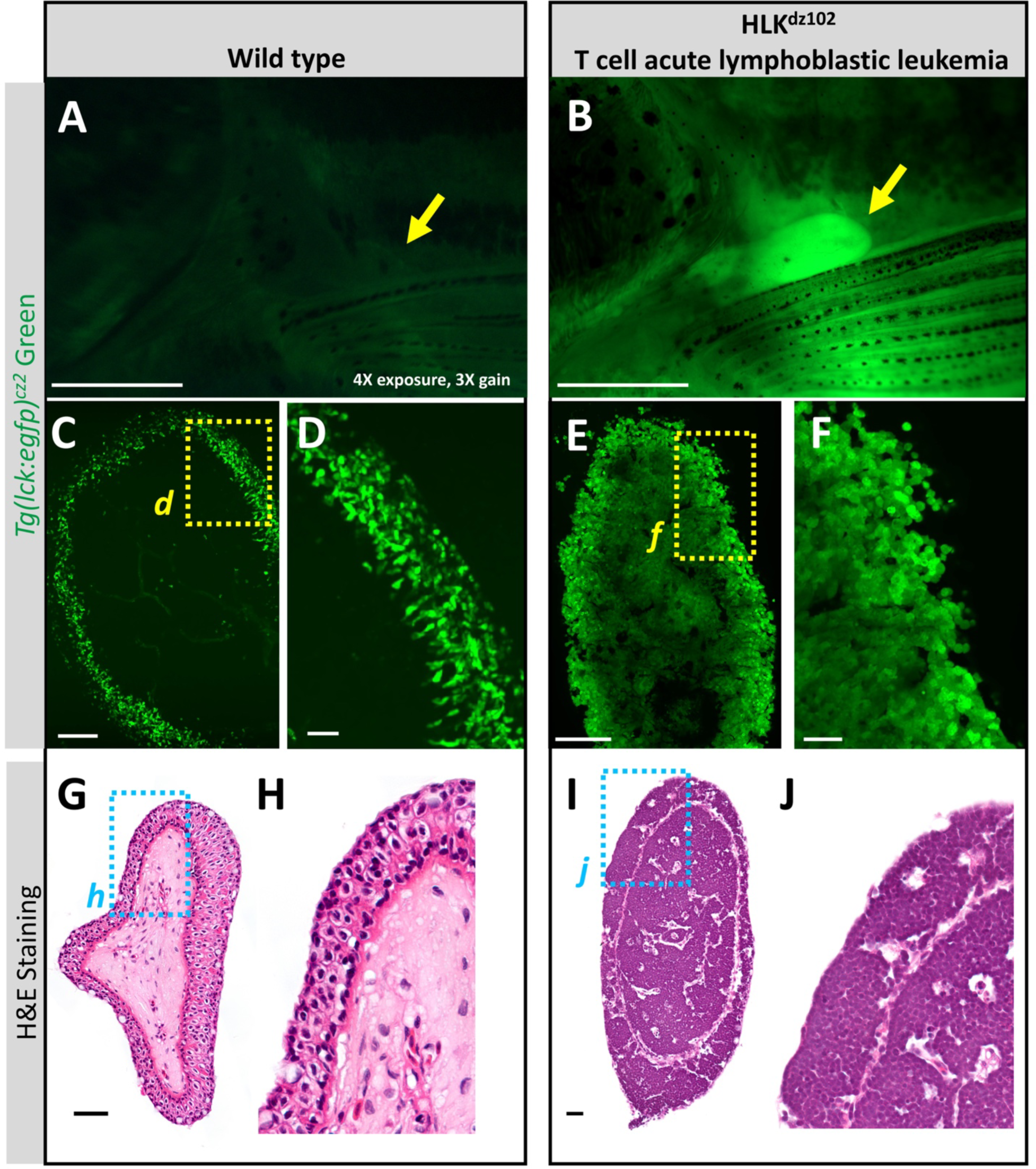
The ALO is heavily infiltrated with leukemic T-cells in a zebrafish model of T-ALL. **A-F.** Fluorescence images of *Tg(lck:EGFP)^cz2^* wild type sibling (A,C,D) or T-ALL HLK*^dz102^* mutant (B,E,F) adult animals. Panels show overview images of the ALO/pectoral area taken with a fluorescent stereomicroscope (A,B), confocal images of most of the ALO (C,E), and higher magnification confocal images of the ALO cortex and underlying medulla (D,F). The images in panels D and F correspond to the boxed regions in panels C and E, respectively. The image in panel A was taken with 4X higher exposure and 3X higher gain than the image in panel B, because this area was much brighter in the T-ALL fish than in the WT fish. **G-J**. H&E stained paraffin histological sections of ALOs from adult wild type (G,H) or T-ALL HLK*^dz102^* mutant (I,J) adult animals. The images in panels H and J correspond to the boxed regions in panels G and I, respectively. The T-ALL HLK*^dz102^* ALO is almost entirely filled with leukemic T cells. Scale bars = 1 mm (A, B), 100 µm (C, E), 25 µm (D,F,G,I).

## Discussion

We have identified a new external organ on adult zebrafish, the axillary lymphoid organ (ALO). The ALO is located behind the operculum of select cyprinids, in the path of water flowing outward from the gills. This novel organ appears in juvenile fish approximately 30 days post-fertilization, is around 10 mm long, and regenerates after being amputated. Histological and ultrastructural examination reveals a stratified cortex with 3 epithelial layers and a reticular medulla containing blood and lymphatic vessels. Single-cell RNA-Seq, *in situ* hybridization, and transgenic reporter lines were used to identify cell types of the ALO. Enormous numbers of immune cells are present in the ALO, and they can be observed trafficking between lymphatic vessels and the ALO cortex using a network of medullary fibroblastic reticular cells as a migration pathway. The basal cortex is densely packed with *lck*^+^ lymphocytes, while B cells and macrophages are more uniformly distributed across the ALO. In animals with T cell leukemias, *lck*^+^ lymphocytes completely infiltrate the cortex and medulla of the ALO. The external location, translucency, and lack of pigment make the ALO ideal for live imaging of immune cells and their surveillance functions.

The small size, translucency, and relatively late appearance in development of the ALO likely contributed to the dearth of previous reports of this organ. On cursory examination it could easily be mistaken for a scale or an isolated fin abnormality. A few previously published reports have used a pectoral fin axial lobe as a taxonomic character for cypriniform fishes (*54, 55*), and other published reports noted the presence of axillary “spines” in other teleosts, including the *Batrachoididae*, but these tissues were not investigated beyond gross anatomy (*56, 57*). External examination of specimens of a small sample of different species in the Smithsonian National Museum of Natural History fish collection suggests that ALOs characterize “basal” teleosts including Danionidae species (**Table S1**). However, the infraclass Teleostei is enormously diverse and species-rich, and we were only able to examine a small, representative sample. Many other teleost fishes likely have ALO-like structures. The fixed teleost specimens in the Smithsonian collection show a diversity of ALO morphologies (**Fig. 1L-O**). Because we did not dissect these specimens as part of this study we could not ascertain the internal morphology of the ALOs of other species or whether similar internal ALO-like structures are present in other fishes. A full review of the ALO throughout teleost fishes would be needed to better understand the taxonomic distribution and range of morphologies of these structures.

The ALO contains enormous numbers of immune cells, especially T and B cells, and our findings suggest that it is a secondary lymphoid organ that may play an important role as a nexus for immune cell trafficking and immune cell signaling. The ALO cortex includes three epithelial cell types that localize to separate cortical layers and that have distinct gene expression profiles and morphological features, such as the characteristic microridges of surface epithelial cells. Basal epithelial cells closest to the ALO medulla express copious transcripts for the chemokine ligands *ccl25a* and *ccl25b*, like gut-associated lymphoid tissue of mammals (**Figure 6B**) (*58*), and appear to be an important signaling hub for immune cell recruitment (**Figure 7C**). This is corroborated by live imaging of immune cell transgenic lines, notably the T lymphocyte-specific *Tg(lck:egfp)^cz2^* reporter line, which shows large numbers of highly active T cells aggregating specifically in the basal most area of the ALO cortex (**Supp. Movie 9**, see 0:18-0:31). The cortical basal epithelial cells lie in close proximity to the sheetlike cell processes and cell bodies of fibroblastic reticular cells (FRCs), located just on the other side of the basal lamina in the adjacent ALO medulla. FRCs are specialized mesodermal cells found in lymph nodes and other mammalian immune organs that have important and well-documented roles in immune cell trafficking in these organs. In zebrafish, FRC-like cells have been identified in larval kidney and hematopoietic tissues, although it is not clear how closely these resemble mammalian FRCs (*43, 59–62*). ALO FRCs express characteristic markers of mammalian lymph node FRCs (**Figure 6A-E**), and they have similar thin sheetlike cell processes with dense collagen-rich matrices. Three-dimensional visualization of the FRC network in the zebrafish ALO from confocal images of *TgBAC(pdgfrb:egfp)^ncv22^* transgenic animals or from array tomography of the ALO (**Supp. Movie 5**) show that the sheetlike interconnected extensions of FRCs form radial highways that emanate from lymphatic vessels, where many FRC cell bodies are located (**Fig. 6K, Supp. Fig. 9, Supp. Movie 5**), outward to the cortex, similar to the three-dimensional networks of FRCs found in mammalian lymph nodes (*63–65*). Like cortical basal epithelial cells, medullary FRCs of the ALO also appear to be major hubs for chemokine signaling to immune cells (**Figure 7C**). Transmission electron micrographs show immune cells closely associating with FRC cell bodies and processes (**Figure 6I,J**), and live imaging reveals multiple types of immune cells actively migrating along FRCs, between the ALO cortex and lymphatic vessels in the ALO medulla although further investigation is needed to determine the nature and function of immune cell migration between the ALO cortex and medullary vessels (**Supp. Movies 6-9**). The lymphatic vessels of the ALO themselves drain to a large adjacent lymph sac (**Fig. 5E-G**), and a dense conduit of T cells can be seen between the nearby thymus and the ALO, especially in juvenile zebrafish (**Fig. 7S,T, Supp. Fig. 10A,B**), connecting the ALO to a recently described path between the sub-operculum and thymus (*11*). Together, the ALO bears many of the hallmark features of a secondary lymphoid organ (SLO), and its highly accessible location makes it ripe for further detailed experimental analysis and comparative morphology.

Secondary immune tissues have been described in several teleost species, although their architecture can differ somewhat from those found in higher vertebrates (*12, 66*). In mammals, SLOs are locations where antigens are concentrated and presented by professional antigen presenters to pools of naïve surveilling lymphocytes. In fishes, lymphocyte activation and affinity maturation of B cells have been characterized in melanomacrophage centers of the head kidney marrow and the spleen, but these tissues are not easily live imaged (*8, 67, 68*). The ALO resembles other teleost mucosa-associated lymphoid tissues (MALTs), including the organized nasal-associated lymphoid tissue (O-NALT), bursa, and gut-associated lymphoid tissue (GALT) where B and T cells tend to be interspersed (*9–11, 13, 69*). The ALO is also likely contiguous with the epidermis, where a tessellated lymphoid network (TLN) was recently described that contains streaming T cells just under the scale junctions (*18*). Plausibly, all MALTs in the adult zebrafish may be interconnected, although extensive additional imaging and experimental analysis would be needed to examine this.

In addition to three epithelial cell populations (surface, mid-level, and basal), our single cell RNA-Seq, *in situ* hybridization, array tomography, and transgenic reporter line characterization shows that the ALO contains several additional cell types in its cortex, some of which may also have accessory roles in immune defense. Like mammals, fishes have sensory cells that detect external stimuli including touch and water-borne chemicals and molecules. The ALO has Merkel cells that transduce touch (*31, 70*) and two solitary chemosensory cell types, tuft-like cells that sense the environment and secrete leukotrienes to communicate with neurons and immune cells (*34*) and taste-like sensory cells (**Fig. 4K-P, Supp. Figs. 5-7, Supp. Movies 2,3**). As noted previously (*31*), Merkel cells lie very close to the ALO surface and extend a single ventral process (**Supp. Fig. 5E,G**). Interestingly, our array tomography data reveals that the solitary chemosensory cells each have two separate ciliary apical extensions protruding out to the external environment (**Supp. Movie 3**), suggesting that individual cells might be able to receive two separate differential external chemosensory signals. Sensory cells such as these could potentially be communicating with immune cells in neuroimmune cellular units (*71*). In mammals, dermal sensory neurons in the skin can change the transcriptional states of immune cells in adjacent patches of skin (*72*), and lymph nodes are innervated and modulated by sensory neurons (*73*). Although it remains unclear whether and to what extent ALO sensory cells interface with immune cells, this organ would provide a superb platform for studying any such interactions *in vivo.* The ALO cortex also includes characteristic mucus-producing goblet cells (**Fig. 4I,J, Supp. Fig. 4, Supp. Movie 3**). These cells produce and secrete mucins in the respiratory, reproductive and gastrointestinal tracts of mammals, and in the gut and skin of teleost fishes, where they play an important role in role in immunity both by creating a passive barrier to infection and by actively participating in immune responses (*74*). Goblet cells in mammals have been shown to actively endocytose soluble substances and pathogens and to transmit these antigens to underlying antigen-presenting cells. Goblet cells in the ALO similarly endocytose and accumulate foreign substances such as bacterial lipopolysaccharide (LPS; **Supp. Fig. 4F-G**), suggesting the ALO may also serve as a useful model for detailed *in vivo* dissection of goblet-immune cell interactions. In addition to goblet cells, the ALO cortex contains abundant large ovoid cells with uniformly sparse cytoplasm and complex folded or multilobed nuclei, characteristic features of club cells (**Supp. Movie 2**). Related to human cells found in the airway epithelium (*35*), in teleosts these cells are thought to have a role both in conspecific fear responses to injured fish, via release of a “fright substance” (“Shreckstoff”) from injured animals, as well as in immune responses, possibly via internalization of bacterial-laden mucus (*37, 38, 75*). Although we were not able to recover a cluster corresponding to these cells in our scRNAseq data, possibly due to their very large size and fragility, we discovered fortuitously that these cells are weakly positive for the same *Tg(gng13a:eGFP)^y709^*transgene that we used to visualize chemosensory type 2 cells (**Fig. 4O-R**), making it possible to observe and study them using confocal imaging. Again, the ALO should provide a valuable platform for *in vivo* investigation of these unusual cells.

In conclusion, we have identified a novel external and highly accessible organ with putative immune surveillance functions in the zebrafish. As a vertebrate species with well-developed genomic resources, a powerful toolkit of methods for imaging of developing and adult animals, and a vast number of transgenic lines available for visualizing almost any cell type of interest *in vivo,* the zebrafish and its axillary lymphoid organ (ALO) will provide a new powerful model for high resolution imaging and experimental manipulation of secondary lymphoid organ function.

## Materials and Methods

### Fish husbandry and fish strains

Fish were housed in a large zebrafish-dedicated recirculating aquaculture facility (4 separate 22,000L systems) in 6L and 1.8L tanks. Fry were fed rotifers and adults were fed Gemma Micro 300 (Skretting) once per day. Water quality parameters were routinely measured, and appropriate measures were taken to maintain water quality stability (water quality data available upon request). The following transgenic and mutant fish lines were used for this study: *Tg(mrc1a:eGFP*)*^y251^* (*40*), *Tg(pdgfrb:eGFP)^ncv22^* (*76*), *Tg(kdrl:mcherry)^y205^* (*41*), *Tg(lyz:DsRed2)^nz50^* (*77*), *Tg(lyve1:DsRed2)^nz101^* (*78*), *Tg(CD79b:EGFP)^fcc89^* (*48*), *Tg(lck:mcherry)^ns107^* (*79*), *Tg(mpeg1:EGFP)^gl22^* (*80*) *Tg(lck:EGFP)*^cz1^, *Tg(lck:EGFP)^cz2^* (*52*)*, Tg(atoh1a:nls-Eos)^w214^* (*32*), *HLK^DZ102/DZ102^* (*81*), and *Tg(gng13a:eGFP)^y709^*. Some fish were maintained and imaged in a *casper* (*roy, nacre* double mutant (*82*)) genetic background to increase clarity for visualization by eliminating melanocyte and iridophore cell populations to prevent them from obscuring images. This study was performed in an AAALAC accredited facility under an active research project overseen by the NICHD ACUC, Animal Study Proposal # 21-015.

### Taxonomic analysis of axillary lymphoid organ phylogeny

Specimens were examined from the formalin-fixed alcohol preserved collections in the Division of Fishes, National Museum of Natural History, Smithsonian Institution to determine presence or absence of the fleshy pectoral lobe. Species were chosen that were closely related to the zebrafish, as well as others in the larger taxon Ostariophysi. The full list of taxa examined for the presence of axillary lymphoid organs is listed in **Table S1**. Specimens were photographed and their size recorded. Images in **Figure 1L-O** show pectoral lobes on representative Danionidae specimens of *Rasbora caverii* and *Laubuka* sp., as well as examples of lobes from outgroup specimens of Atlantic tarpon (*Megalops atlanticus*) and Milkfish (*Chanos chanos*).

### Histology and transmission electron microscopy

Adult zebrafish were euthanized by placing them in an ice bath for 10 minutes. For histology, the caudal 1/3 of the animal was removed to improve fixation penetration and the sample was placed in 4% paraformaldehyde shaking overnight. The following day, the fish were rinsed 3 times with phosphate buffer saline and transferred to 70% ethanol. Samples were then sent to Histoserv, Inc. (histoservinc.com) for sectioning and histological staining. Slides were imaged on a Leica DMI 6000B inverted compound microscope with Leica DMC6200 camera, Leica Application Suite X software, and 10X 0.3 NA air, 20X 0.8 NA air, 40X 1.25 NA oil, and 63X 1.32 NA oil Leica objective lenses.

For transmission electron microscopy zebrafish ALOs were removed post euthanasia and fixed with 4% glutaraldehyde in 0.1M sodium cacodylate buffer, pH 7.4. After fixation samples were inserted into mPrep tissue capsules and loaded onto an mPrep ASP-2000 Automated Biological Specimen Preparation Processor (Microscopy Innovations, LLC, Marshfield, WI.) which automates all subsequent processing steps including the following: post-fixation in 2% osmium tetroxide, en-bloc in 2% uranyl acetate (aqueous), dehydrated in a graded ethanol series followed by further dehydration in 100% acetone, and finally infiltrated and embedded in Embed 812 epoxy resin (Electron Microscopy Sciences Hatfield, PA.). Embedded samples were polymerized in an oven set at 60°C. Samples were then ultra-thin sectioned (90nm) on a Leica EM UC7 Ultramicrotome. Thin sections were picked up and placed on 200 mesh cooper grids and post-stained with UranyLess (Uranyl Acetate substitute, Electron Microscopy Sciences, Hatfield, PA.). and lead citrate. Imaging was performed on a JEOL-1400 Transmission Electron Microscope operating at 80kV and an AMT BioSprint-29 camera.

### Array Tomography

ALOs were amputated (described below) from wild type EK strain zebrafish. Whole ALOs were fixed with Karnovsky’s fixative and post-fixed in 2% OsO_4_ and 1.5% potassium ferricyanide in 0.1 M sodium cacodylate for 1 hr at room temperature (RT). Next, whole ALOs were washed with ultrapure water and stained with 1% aqueous uranyl acetate for 1 hr at RT. After washing with water, the ALOs were treated with lead aspartate at 60°C for 30 minutes and washed again with water. The ALOs were dehydrated through a graded ethanol series (10 min. each of 35%, 50%, 70%, 95%, and 100% x 3) and finally in 100% propylene oxide (PO). After dehydration, the ALOs were infiltrated with increasing amounts of polybed resin with PO (resin: PO; 1:3, 1:1, 3:1, and 100% resin). Finally, the samples were placed in 100% degassed resin in flat beam capsules (EMS), and cured at 65°C for 48 hrs. After polymerization, resin pieces containing an ALO were trimmed with a Leica EM TRIM2 milling system.

After trimming the block face down, 100nm serial sections were collected with a 45° ultra-diamond knife (Diatome) using an ATUMtome (RMC Boeckeler) on carbon nano tube PEN tape (Teijin). Tape strips were mounted onto 12 mm double sided carbon tape with aluminum base (EMS) and attached to 4-inch type-p silicon wafers (EMS) and grounded with conductive graphene carbon paint (EMS). Overview wafer images were collected using a Canon EOS 1300D camera. Wafers were affixed to a 4-inch stage-decel holder in a Zeiss SEM (GeminiSEM 450; Carl Zeiss) and imaged using ATLAS 5 Array Tomography software (Fibics). Five wafers were imaged using a four-quadrant backscatter detector, with the electron beam operated at 3 kV EHT with 1 kV beam deceleration and 800pA probe current. Low-resolution overview scans were collected at 3000 nm pixel resolution, medium-resolution section sets were collected at 150 nm pixel resolution, and high-resolution sites were collected at 25 and 10 nm pixel resolution. Once image acquisition was complete, the image stack was locally cropped and aligned using the ATLAS 5 software. The resulting image stack was exported and then processed using python-based scripts to produce two aligned and contrast/brightness adjusted image datasets: a stack of images at 10 or 25 nm (xy) x 100 nm (z), and a final binned 100×100×100nm isotropic volume. Raw array tomography image data is deposited at EMPIAR.

### ALO amputation and *ex vivo* imaging

Adult zebrafish were anesthetized in buffered tricaine (160 mg/L) and placed on a moist paper towel under a dissecting microscope (Leica M165) with goose neck illumination. Using #55 sharp forceps (Dumont 11255-20) to grab the base of the ALO, the ALO was removed with a gentle tug of the forceps. The amputated ALO was then placed in a MatTek dish (# P35-1.5-14-C) containing 100-150 µl of 10% fetal bovine serum in PBS. A 25 mm round coverslip was then placed on top of the solution containing the ALO. A similar setup was used for ex vivo imaging on our upright microscope, except we used a 50 mm dish (MatTek P50G-1.5-30-F) with a 35 mm coverslip (MatTek PCS-1.5-35-NON), to account for the large diameter of the 20X 1.0 NA water immersion objective.

### Live staining and *in vivo* ALO treatments

Adult and subadult zebrafish were stained with BODIPY membrane stain to visualize external structures by soaking them in 100 ml of 20 ng/ml BODIPY^TM^ 630/650 (ThermoFisher Cat # D10000) in aquaria system water for one hour. After soaking, fish were transferred to fresh aquaria system water before anesthesia and live imaging. Lipopolysaccharide (LPS) treatment was carried out by anesthetizing adult zebrafish in 160 mg/L Tricaine solution and placing them on a moist paper towel under a dissecting microscope, then placing 25 µl of LPS solution (1mg/ml Cy5 labeled, Nanocs Cat # LPS-S5-1) directly on the ALO. The fish was then flipped over, and the process was repeated on the other ALO. A kimwipe (Kimtech Science 05511) wetted with anesthesia water was placed on top of the fish, and allowed to sit for three minutes. The fish was transferred to fresh system water for recovery until fully revived, (about two minutes) then re-anesthetized followed immediately by removal of the ALOs for imaging.

### Angiography and lymphatic drainage

ALO lymphatic drainage was assessed by anesthetizing fish in 160 mg/ml tricaine, placing them in a slit cut into a wet sponge, then injecting 50 nl of undiluted (2 uM) Qtracker^TM^ 705 Vascular labels (Invitrogen cat# Q21061MP) directly into the ALO using a Drummond Nanoject III microinjector (Item# 3-000-207) with pulled glass capillary needles (Drummond item # 3-00-203-G/X). Fish were then placed back into anesthesia solution and then moved to an imaging dish (Lab-TekII #155360) with tricaine solution and gently covered with a sponge to prevent movement. Flow though ALO blood vessels was assessed by using angiographic injection of Hoechst 33342 dye to label the nuclei of circulating red blood cells. Injections were performed as noted above but with 250 nl of Hoechst 33342 (1 µM solution Cat # H3570 diluted 1:1 with PBS), and with injection into the caudal axial vasculature instead of into the ALO. Fish were placed in fresh aquaria system water for recovery until fully revived (about two minutes), then re-anesthetized and imaged.

### Image Acquisition

Confocal images were acquired using either a Nikon Ti2 inverted microscope with Yokogawa CSU-W1 spinning disk confocal (Hamamatsu Orca Fusion-BT camera), a Nikon Ti2 inverted microscope with Nikon A1R scanning confocal, or a Nikon FNSP upright microscope with AXR scanning confocal. 405 nm, 488 nm, 561 nm, 640 nm laser lines were used on all systems. The following Nikon objectives were used: 4X Air 0.2 N.A., 10X Air 0.45 N.A., 20X water immersion 0.95 N.A., 20X water immersion 1.0 N.A., 40X water immersion 1.15 NA. Stereo microscope pictures were taken using a Leica M165 microscope with Leica DFC 7000T camera and Leica Application Suite X software.

### Image Processing

Images obtained from the Nikon A1 and AXR resonant scanners were denoised using Nikon Denoise AI. A median filter with a kernel size of 3 was applied to some images. A subset of images were deconvolved using NIS Batch deconvolution. Images were processed using NIS Elements. Maximum intensity projections of confocal stacks are shown for fluorescent confocal images. When fluorescent and DIC images are shown together, extended depth of focus images of stacks are shown. When needed, timelapse movies were aligned in XY using NIS Elements. Non-linear adjustments (gamma) were made to some images to improve the visualization of images with high dynamic range. Images adjusted: Fig 7 D-O, and Supp Movies 7-9. Time-lapse movies were made using NIS Elements and exported to Adobe Premiere Pro CC 2024. Adobe Premiere Pro CC 2024 and Adobe Photoshop CC 2024 were used to add labels and arrows to movies. Schematics were made using Adobe Photoshop CC 2024, Microsoft PowerPoint, and Bio Render software.

### Preparation of ALO single cell suspension for scRNA-Seq

Adult zebrafish were anesthetized with MS-222 prior to removal of the ALOs. ALOs from 4 different *Tg(lck:EGFP*^cz1^; *lyz:DsRed2^nz50^*double transgenic fish (2 males and 2 females) were removed with forceps and placed in dissociation media (1:1 DMEM:Liberase) (Gibco A1896701/ Sigma-Aldrich 05401119001). The tissue was slowly pipetted up and down at room temperature for 45 minutes with a 1000 µL pipette set to 500 µL to ensure gentle dissociation. Cell dissociation was stopped by adding an equal volume of STOP solution (5% FBS, 2% BSA in DMEM)(Corning 35-010-CV/Millipore Sigma A9418-5G/Gibco A1896701) and gently inverting the tube. Cell suspensions were filtered through a 40-micron filter and spun down at 500 x g for 5 minutes at room temperature. The supernatant was removed, and cells were resuspended in a 1X PBS 2% BSA (Gibco 10010023/Millipore Sigma A9418-5G) solution. Cells were counted on a LUNA-FL (Logos Biosystems) and diluted to the optimum concentration of 700 cells/ml using the same 1X PBS 2% BSA solution. 10,000 cells were loaded onto the 10x Genomics Chromium X controller.

### Sequence alignment and quality control

Alignment of sequencing reads and processing into a digital gene expression matrix was performed using Cell Ranger version 7.0.0. The data was aligned against GRCz11 release 99 (January 2020) using the Lawson Lab Zebrafish Transcriptome Annotation version 4.3.2 (*83*), available from https://www.umassmed.edu/lawson-lab/reagents/zebrafish-transcriptome/. Cells were processed and analyzed using Seurat version 4.3.0.1 (*84*) and R version 4.2.3. Cells with abnormally high (> 2500) or low (< 200) numbers of detected features, or with abnormally high mitochondrial content (> 5%) were removed. The remaining cells were normalized to 10,000 transcripts per cell and scaled using the ScaleData function of Seurat with default settings. Following initial clustering, it was noted that one cluster appeared to group with every other cluster instead of forming a unique identity, and it contained an abnormally high percentage of ribosomal protein genes. The cells in this cluster were removed and the remainder of the dataset was used for all subsequent analyses shown. **Figure 3B** provides the cell and read count metrics before and after performing quality control.

### Dimensionality reduction, clustering, and visualization

2000 variable features were identified for our sample using Seurat’s FindVariableFeatures function with default parameters. Principal component analysis (PCA) was performed with RunPCA algorithm using the determined most variable features. To identify the number of significant PCs for downstream analyses, the JackStrawPlot function of Seurat was used. The FindNeighbors and FindClusters functions from Seurat were applied utilizing the number of significant PCs, to identify clusters. After exploring a variety of possible resolutions 0.15 was selected, generating 14 distinct clusters. A Uniform Manifold Approximation and Projection (UMAP) was calculated using the RunUMAP function and visualized using DimPlot. CellChat v2 (*85*), an open source R package, was used to produce a cellular communication circle plot inferring chemokine signaling pathways within the dataset.

### Differential expression analysis

To identify differentially expressed genes between cell types, we used a negative binomial model as implemented in the Seurat FindAllMarkers function, with a log fold change cutoff of 0.25. Genes were considered differentially expressed if the adjusted *P* value was lower than 0.01. A table of the resulting genes with the highest expression values in each cluster can be found at https://github.com/nichd-Weinstein/Axillary-Lymphoid-Organ.

### Defining cell types

Each of the 14 clusters was manually annotated based on an extensive survey of well-known tissue- and cell type–specific markers. These markers were identified through a variety of databases (Human Protein Atlas, The Zebrafish Information Network, and Daniocell) and through an extensive literature search. Our search and identification were guided by preliminary confocal imaging and electron microscopy of the ALO. For each cell type, at least five marker genes were identified.

### Hybridization Chain Reaction (HCR)

HCR was performed to visualize and confirm the identities of each cell type/cluster. HCR probesets were designed by Molecular Instruments (Molecular Instruments, Los Angeles, CA). ALOs were dissected and fixed in 4% PFA (Electron Microscopy Sciences Cat # 15710) for 2 hours at room temperature, washed three times for 5 minutes each time with 1mL of PBST (1x phosphate-buffered saline and tween20)(Gibco 10010023/Millipore Sigma 9005-64-5) and treated with Proteinase K solution (Thermo Fisher 50 µg/mL in PBST) 10 minutes at room temperature. The samples were washed again twice with PBST (5 mins each time at room temperature), before being postfixed with 4% PFA for 20 minutes at room temperature. This was followed by another PBST wash cycle (three times for 5 minutes each time). Fixed ALOs were pre-hybridized with preheated (37°C) HCR probe hybridization buffer (Molecular instruments, HCR™ RNA-FISH Bundle) for 30 minutes at 37°C, rotating at 30 rpm. Hybridization was performed with 2 μL of each 1 μM probe diluted in 500 μL of probe hybridization buffer at 37°C, rotating at 30 rpm, for 12-16 hours. Probe solution was removed by washing with preheated HCR probe wash buffer (Molecular instruments, HCR™ RNA-FISH Bundle) four times 15 minutes each at 37°C, followed by two 5 minute washes with 5X SSCT (12.5 ml 20X SSC in 50 µL tween 20)(KD Medical Cat # RGF-3240/Millipore Sigma 9005-64-5) at room temperature before pre-amplification stage. ALOs were pre-amplified with HCR probe amplification buffer (Molecular instruments, HCR™ RNA-FISH Bundle) for 30 minutes at room temperature. Hairpins (10μL of 3μM stock) were pre-annealed (95°C for 90 seconds and 25°C for 30 minutes with a ramp rate of −0.1°C per second) to create hairpin secondary structure. Hairpins were then mixed with 500μL of fresh HCR probe amplification buffer. The pre-amplification buffer was removed from the ALOs before the addition of the newly mixed Hairpin solution, followed by an incubation period for 12-16 hours in the dark at room temperature. Excess hairpins were then removed and samples were washed five times with 5X SSCT at room temperature and stored at 4°C in the dark until imaging.

## Acknowledgments

The authors would like to thank members of the Weinstein and Sheppard laboratories for their critical comments on this manuscript. The Authors would also like to thank the Research Animal Branch of the *Eunice Kennedy Shriver* National Institute of Child Health and Human Development as well as the RAMB contract animal management staff for excellent animal care and husbandry. We also thank Irene Salinas and Benjamin Garcia for helpful discussions.

## Funding

NICHD, NIH intramural support ZIA-HD008915 (BMW)

NICHD, NIH intramural support ZIA-HD008808 (BMW)

NICHD, NIH intramural support ZIA-HD001011 (BMW)

NCI, NIH Contract No. 75N91019D00024 (KN)

The Herbert R. and Evelyn Axelrod Endowment, Division of Fishes, National Museum of Natural History (DNL, LRP)

Hyundai Hope On Wheels Hope Scholar award (JKF)

OUHSC Stephenson Cancer Center Pilot Grant (JKF)

Oklahoma Center for Adult Stem Cell Research (JKF)

Canadian Institutes of Health Research MOP77746 (EF)

R35 GM118027 (AH)

F32 GM146398 (TFR)

## Author contributions

Conceptualization: DC, MIK, AK, BMW

Methodology: DC, MIK, AK, CD, JSP, GP, DNL, LRP, JKF, KN, BMW

Investigation: DC, MIK, AK, CD, JSP, MVG, GP, DNL, LD, MM, JI, KT, VP, RJW, LB,TFR, YH

Visualization: DC, MIK, AK, DNL, BMW

Supervision: AH, EF, LRP, JKF, KN, BMW

Writing—original draft: DC, MIK, AK, BMW

Writing—review & editing: DC, MIK, AK, CD, JSP, MVG, GP, DNL, LD, MM, JI, KT, VP, RJW, LB, TFR, YH, AH, EF, LRP, JKF, KN, BMW

## Competing interests

Authors declare that they have no competing interests.

## Data and materials availability

Raw and processed 10X scRNA-seq data can be found on GEO using accession number GSE270797 The code used to analyze and visualize the data can be found at https://github.com/nichd-Weinstein/Axillary-Lymphoid-Organ

Array Tomography data is available at https://www.ebi.ac.uk/empiar/

## Notes

https://github.com/nichd-Weinstein/Axillary-Lymphoid-Organ

### Competing Interest Statement

The authors have declared no competing interest.

