## Supplemental Figures for "The axillary lymphoid organ - an external, experimentally accessible immune organ in the zebrafish"

Daniel Castranova *et al.*

**This PDF file includes:**

Figs. S1 to S10
Table S1
Movies S1 to S10

**Other Supplementary Materials for this manuscript include the following:**
Movies S1 to S10

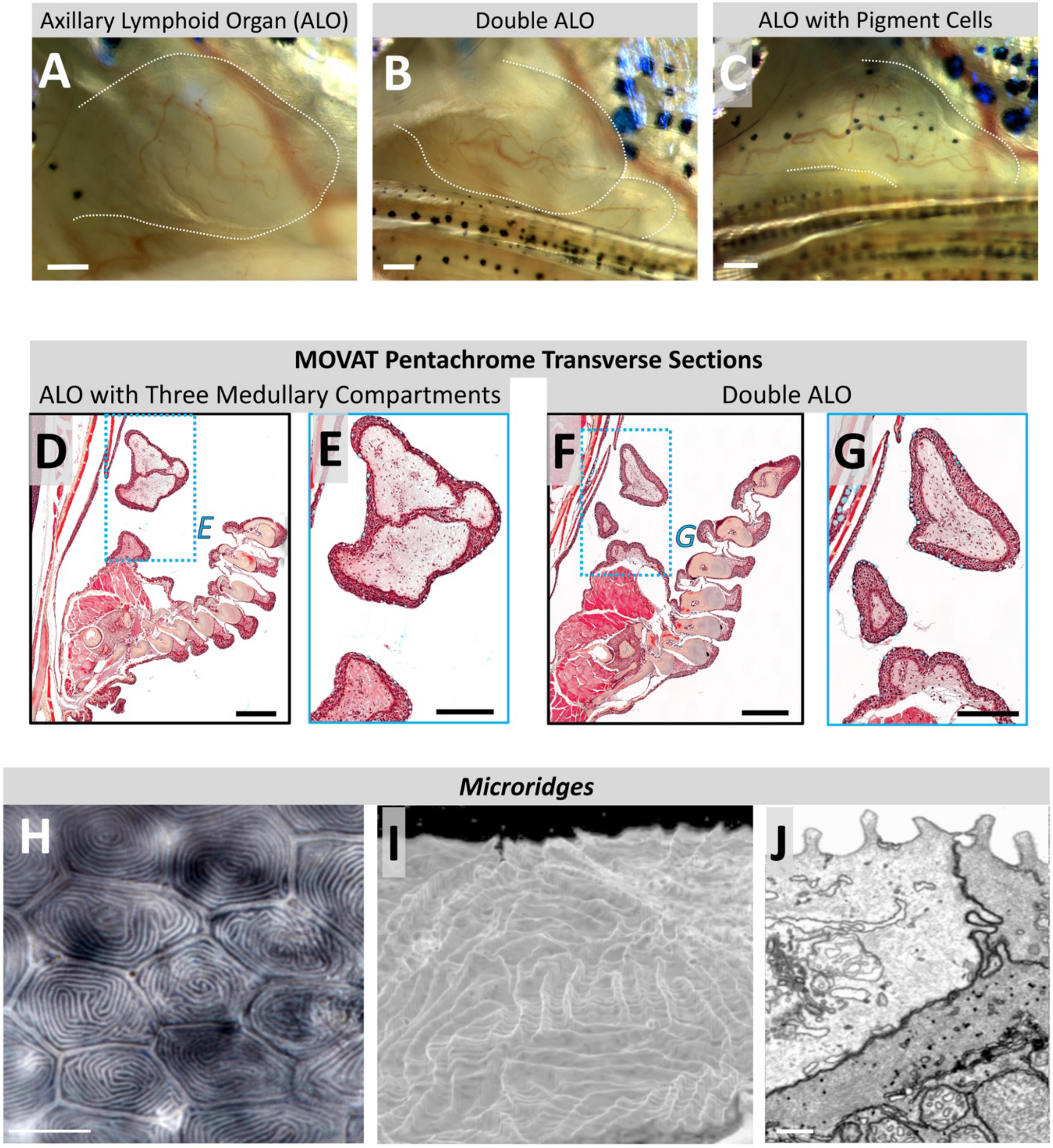


**Fig. S1. Zebrafish axillary lymphoid organ (ALO) variability and dermal microridges**
**A-C.** Stereomicroscopic images of ALOs from different 10 month old EK wild type zebrafish, with dashed white lines indicating the ALO border, showing a single ALO (A), double ALO (B), and an ALO with melanophores on it (C). **D-G.** MOVAT pentachrome stained transverse paraffin sections through the pectoral ALOs of two different adult zebrafish showing their morphological variability, including (D,E) a medulla divided into three compartments and (F,G) a fish with two separated ALOs. The blue dashed boxes in panels D and F show the areas magnified in panels E and G, respectively. **H-J.** Skin microridges on the outside of the ALO shown in a DIC micrograph (H) and in a rendering of an array tomography 3D image volume reconstruction (I,J), with a 3D reconstruction of the entire array in panel I and part of a single plane shown in panel J (see also **Supp. Movies 2 & 3**). Scale bars = 150 µm (A,B,C,E,G), 250 µm (D,F), 10 µm (H), 1 µm (J).

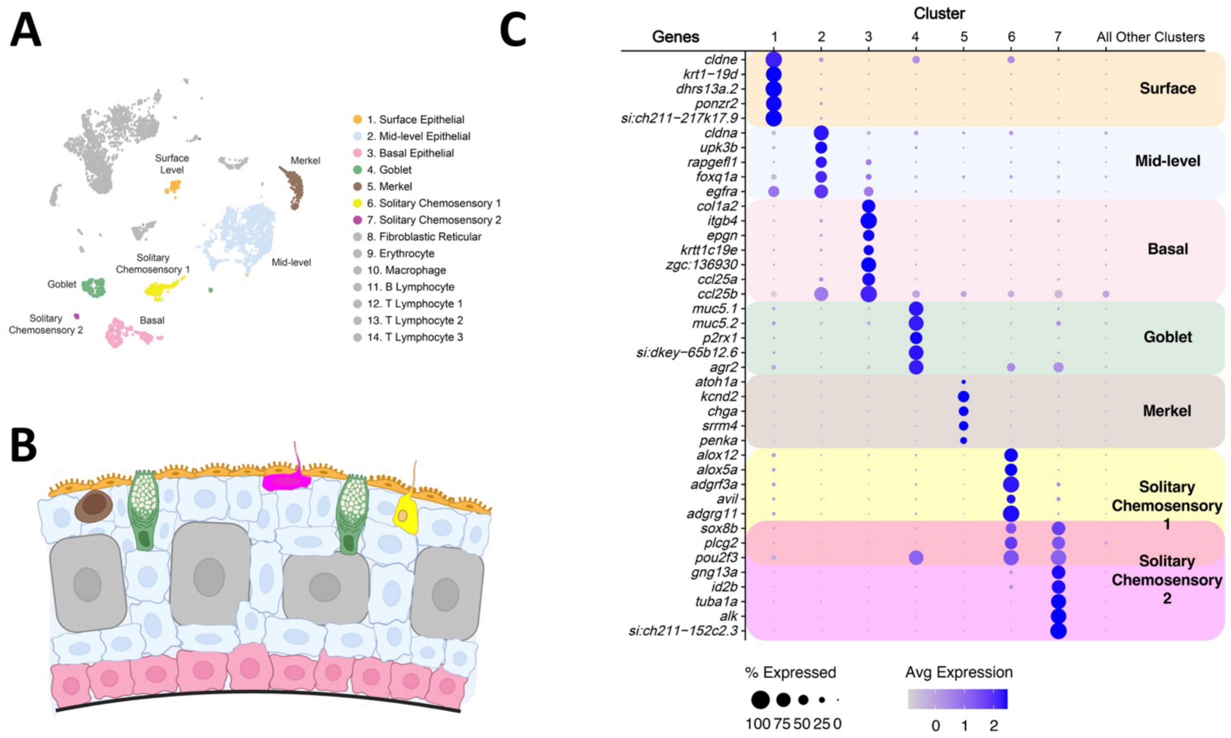


**Fig. S2. Single-cell RNAseq of the ALO cortex**

**A.** UMAP plot of ALO scRNA-seq data highlighting seven clusters that include resident cell types of the ALO cortex. **B**. Schematic diagram of the ALO cortex with the cell types represented by each of the highlighted clusters in panel (A) shown using the same colors. **C.** Dot plot showing the relative expression of genes used to identify and characterize clusters corresponding to resident cell types of the ALO cortex.

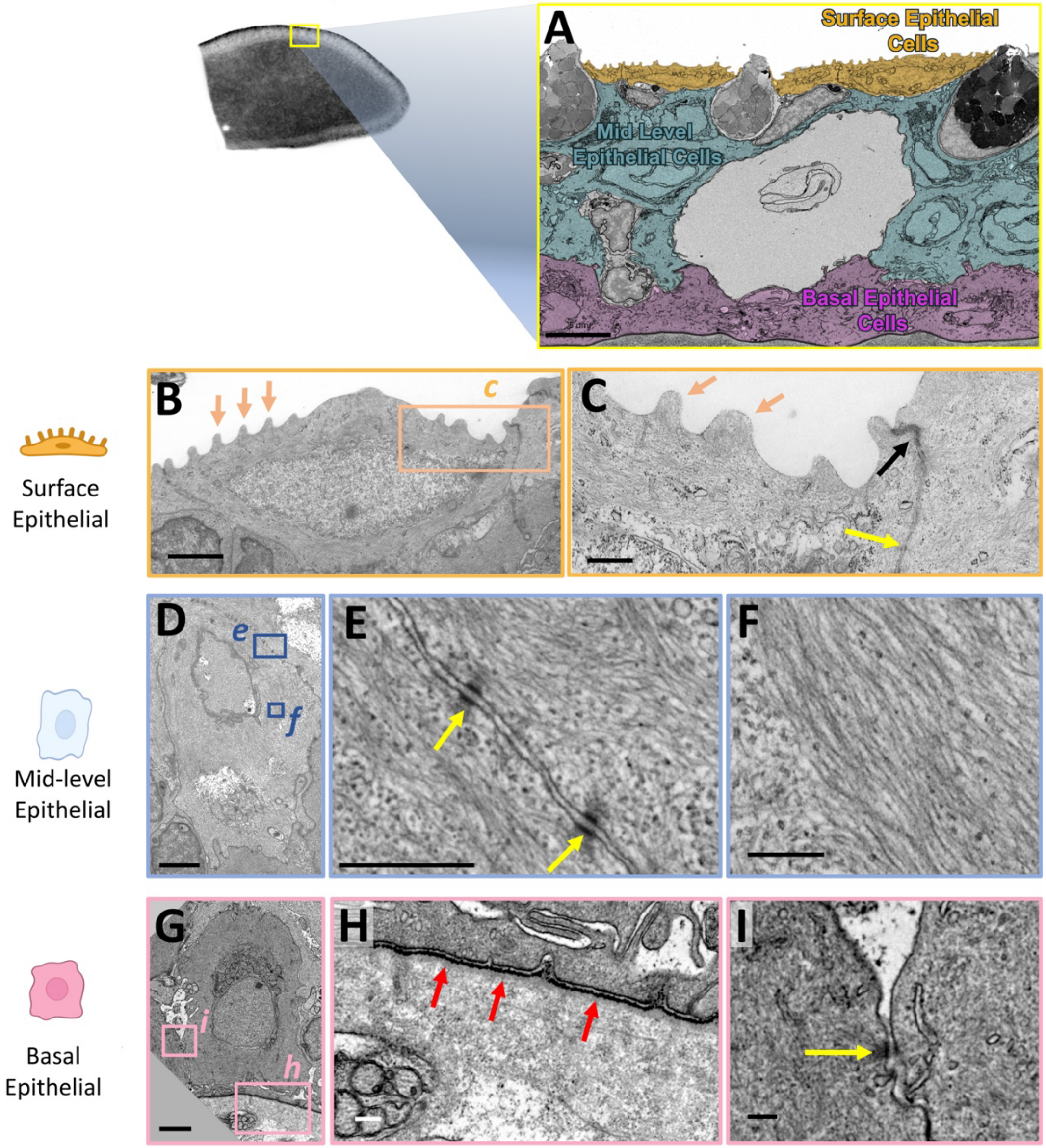


**Fig. S3. Axillary lymphoid organ (ALO) cortical epithelial cells**

**A.** A single pseudocolored section an array tomography image stack of an adult zebrafish pectoral axillary ALO, highlighting the surface, mid-level, and basal epithelial layers. **B,C.** Transmission electron micrograph of a microridge (orange arrows) containing surface epithelial cell. Panel C shows a higher magnification image of the orange boxed area in panel B, with a cell-cell tight junction (black arrow) and desmosome (yellow arrow) noted. **D-F.** Transmission electron micrograph of a mid-level epithelial cell. Panels E and F show higher magnification images of the blue boxed areas in panel D, with desmosomes (yellow arrows in panel E) and intermediate filaments (panel F) noted. **G-I.** Transmission electron micrograph of a basal epithelial cell. Panels H and I show higher magnification images of the pink boxed areas in panel G, with the basement membrane separating the dermal cortex and the medulla (red arrows in panel H) and a basal cell-basal cell desmosome (panel J) noted. See **Supp. Movie 2** for additional array tomography images of cortical epithelial cells. Scale bars = 5 µm (A), 2 µm (B,D,G), 500 nm (C,E,H), 200 nm Type or paste caption here. Create a page break and paste in the Figure above the caption.

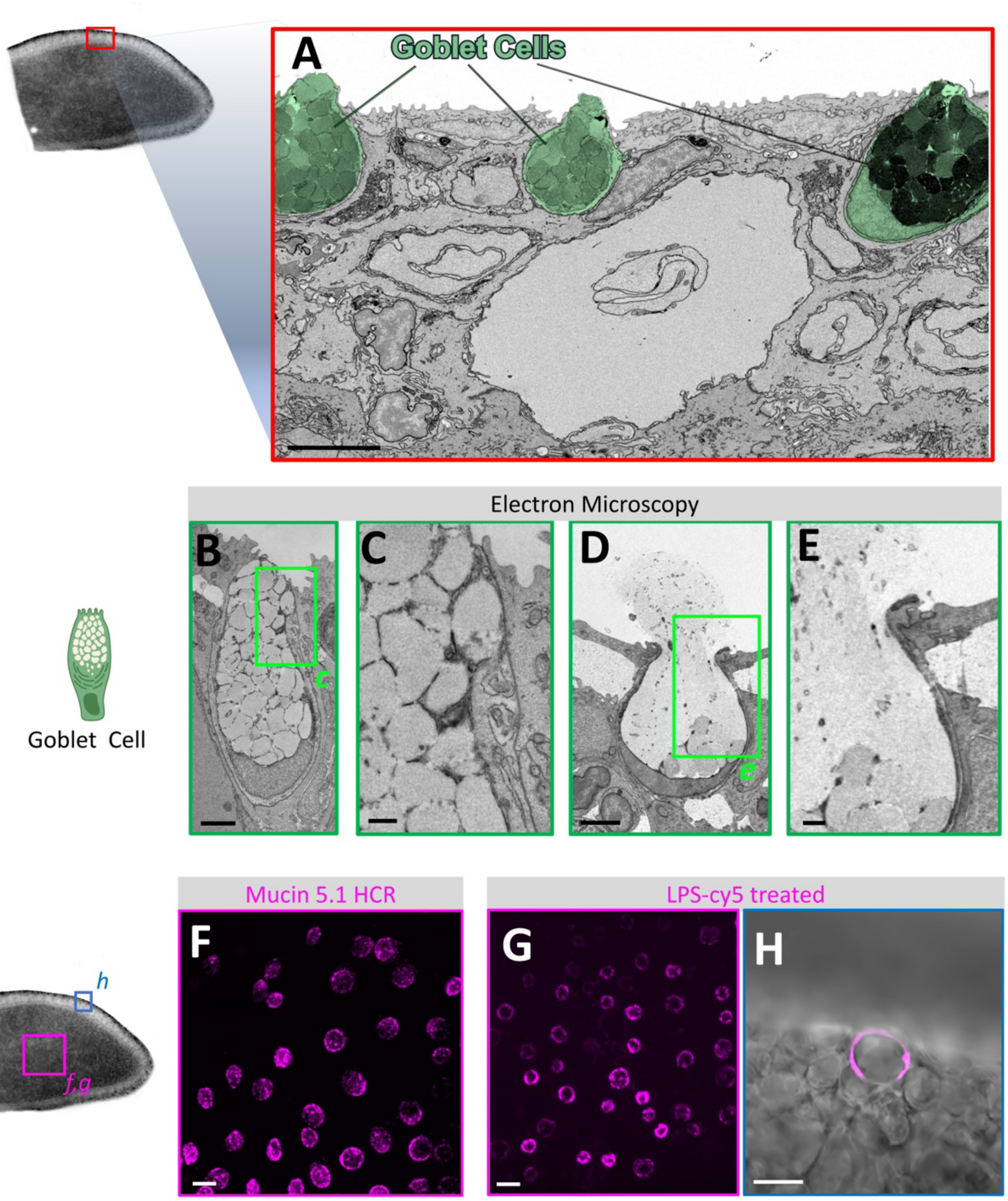


**Fig. S4. Axillary lymphoid organ (ALO) cortical goblet cells**

**A.** A single section from an array tomography image stack of an adult zebrafish pectoral axillary ALO, with goblet cells pseudocolored green. **B-E.** Transmission electron micrographs of goblet cells with intact mucus granules (B,C) or with mucus granules breaking down and mucus being extruded (D,E). Panels C and E show higher magnification images of the green boxed areas in panels B and D, respectively. **F.** Confocal micrograph maximum intensity projection showing hybridization chain reaction (HCR) *in situ* hybridization of an adult ALO probed for mucin 5., showing specific labeling of goblet cells. **G,H.** Confocal micrographs of ALOs treated *in situ* with cy5-labeled lipopolysaccharide (LPS-cy5) and then excised and imaged *ex vivo*, showing specific uptake of LPS-cy5 by goblet cells. Images include a maximum intensity projection (G; comparable view to the image shown in panel F) and a single confocal section with DIC showing a side-view of the dermal cortex with a single LPS-cy5 labeled goblet cell (H). See **Supp. Movies 2 & 3** for additional array tomography images of goblet cells. Scale bars = 5 µm (A), 2 µm (B,D), 500 nm (C,E), 10 µm (F-H).

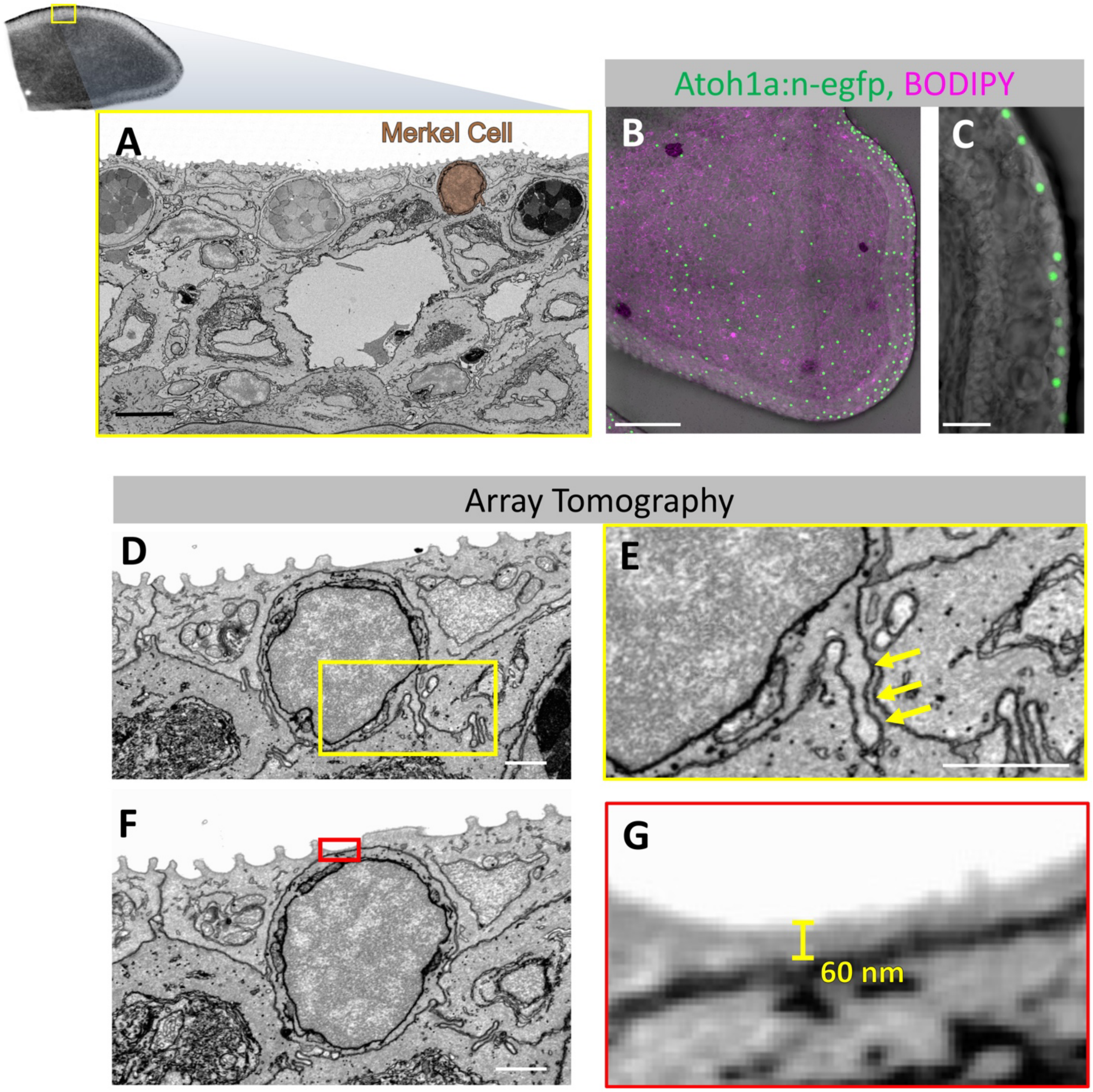


**Fig. S5. Axillary lymphoid organ (ALO) cortical Merkel cells**

**A**. A single section from an array tomography image stack of an adult zebrafish pectoral axillary ALO, with a Merkel cell pseudocolored brown. **B-C.** Confocal/DIC extended depth of focus images of an adult *Tg(atoh1a:nls-Eos)*$^{w214}$ transgenic zebrafish ALO showing Merkel cell nuclei in green. Images include (B) a ALO overview (also with BODIPY 633 in magenta), and (C) a side view of the dermal cortex. **D-G.** Individual sections from transmission electron microscopic array tomography of an adult zebrafish pectoral axillary ALO, showing Merkel cells with large nuclei and small amounts of cytoplasm. Panels E and G show higher magnification images of the yellow and red boxed areas in panels D and F, respectively. The single inwardly projecting microvillus characteristic of Merkel cells is shown in panel E (yellow arrows). The magnified image in panel G shows that the Merkel cell outer membrane is only 60 nm from the outside surface of the ALO. See **Supp. Movie 2** for additional array tomography images of Merkel cells. Scale bars = 5 µm (A), 100 µm (B), 20 µm (C), 1 µm (D-F), 60 nm (G).

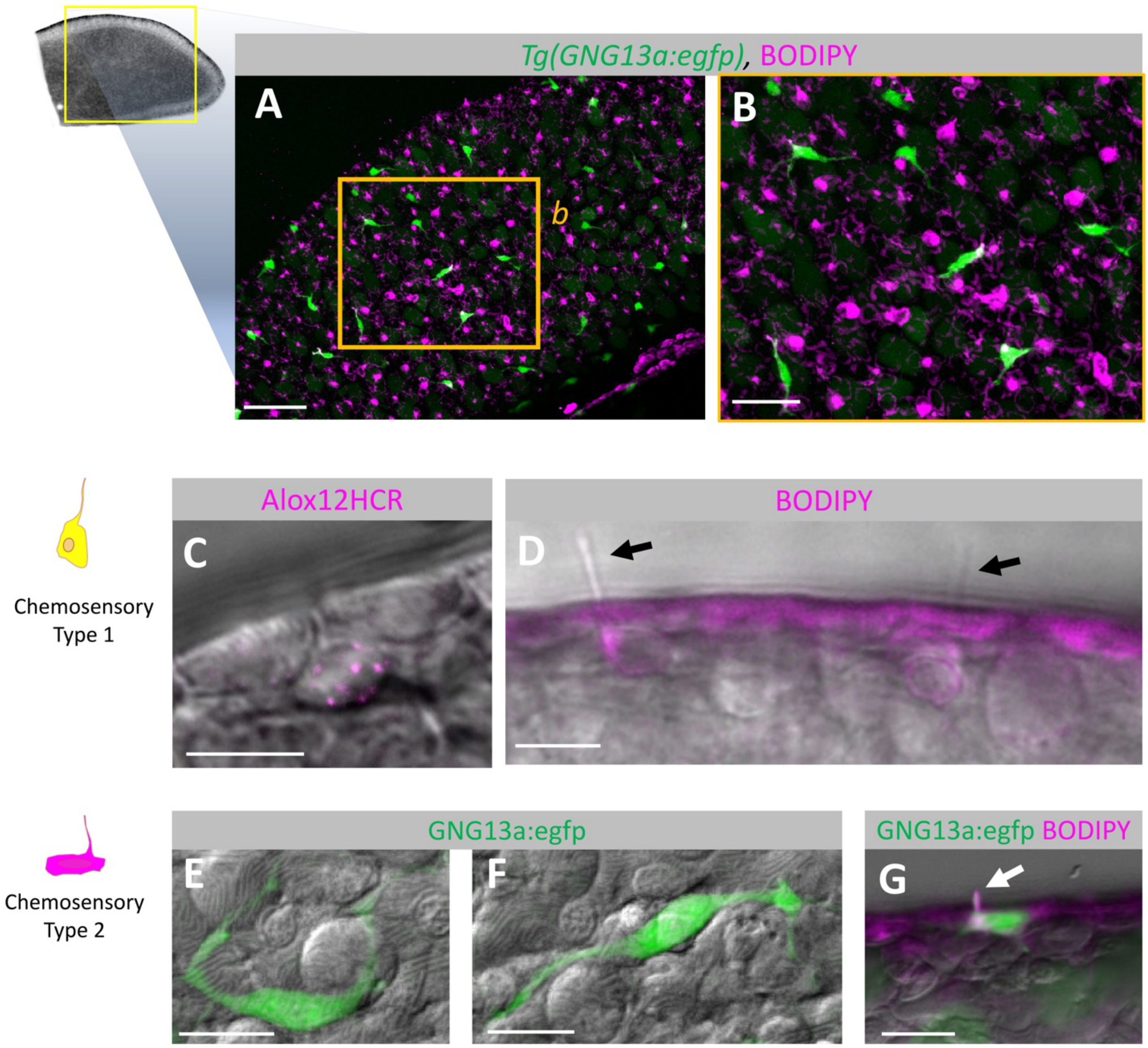


**Fig. S6. The axillary lymphoid organ (ALO) contains two different types of chemosensory cells**
**A-B**. Confocal micrograph maximum intensity projections of a ALO from a *Tg(gng13a:eGFP)*$^{y709}$ transgenic zebrafish treated with BODIPY 633, showing uptake of the fluorescent dye by chemosensory cells. Panel B shows a higher magnification image of the orange boxed area in panel A, with a (**A**) and a close-up of the orange boxed area in (**B**) showing only a subset of chemosensory cells are gng13a positive. **C,D**. Single-plane side-view confocal/DIC micrographs of the cortical surface of wild type adult zebrafish ALOs either (C) probed for HCR with *alox12* showing expression in a chemosensory cell located just below two surface epithelial cells, or (D) treated with BODIPY 633 showing uptake by chemosensory cells, as well as chemosensory cell microvilli extending into the environment (black arrows). **E-G.** Confocal micrograph extended depth of focus projections of ALOs from a *Tg(gng13a:eGFP)*$^{y709}$ transgenic adult zebrafish, showing (E,F) overview images of EGFP-positive chemosensory cells with long projections just underneath the surface of the ALO cortex, and (G) a side view of the ALO cortex of a fish treated with BODIPY 633, showing uptake by the chemosensory cell microvillus (white arrow) and cell body. Scale bars = 50 µm (A), 25 µm (B), 10 µm (C-G)

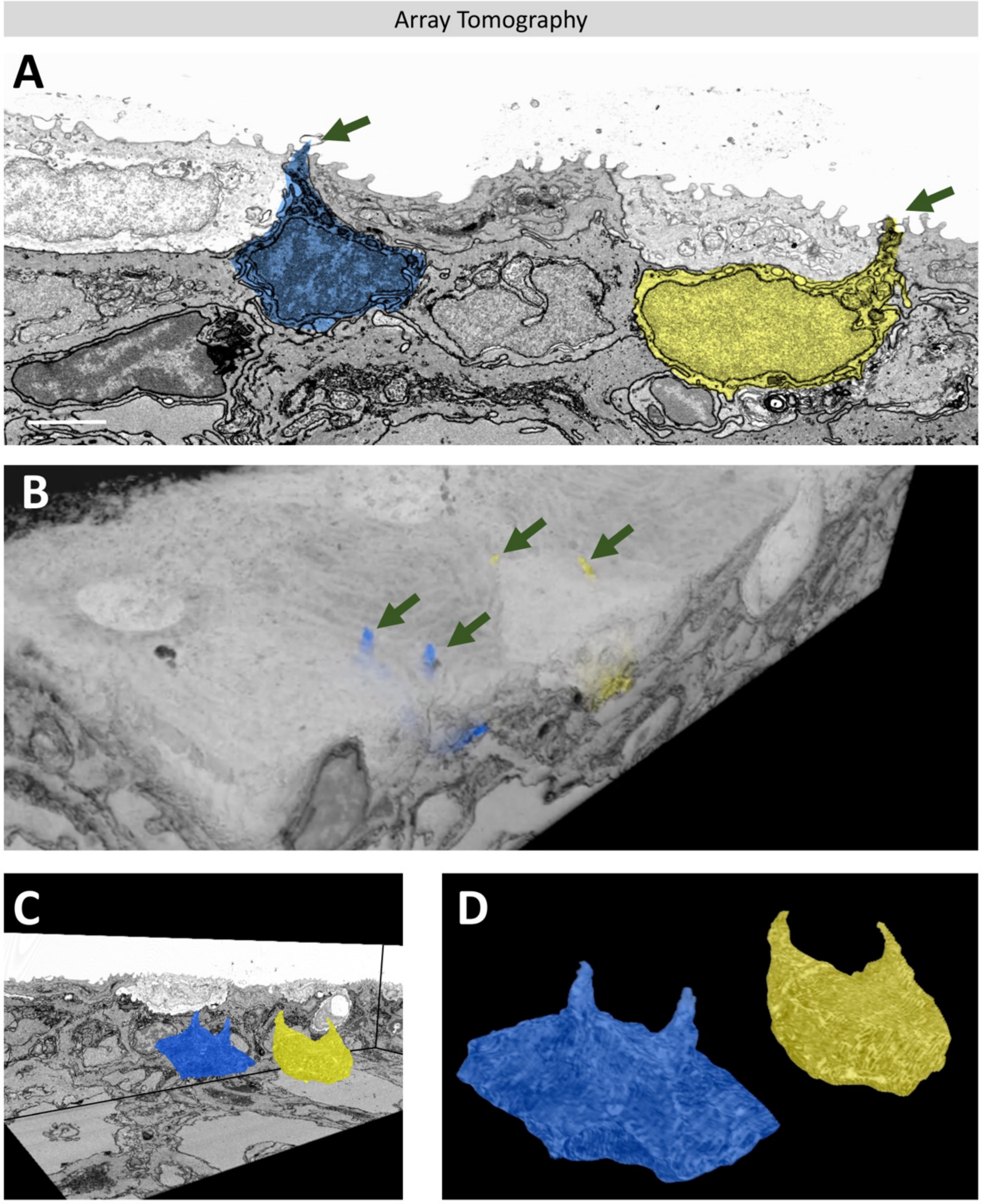


**Fig. S7. ALO chemosensory cells have two external microvilli**

**A.** A single section from an array tomography image stack of an adult zebrafish pectoral axillary ALO cortex, with two separate chemosensory cells pseudocolored blue and yellow. Single sections such as these show individual chemosensory cell microvilli extending from these cells into the environment (green arrows). **B-D.** Volume reconstructions of the same array tomography data set of an adult zebrafish pectoral axillary ALO cortex from which the image in panel A was taken, with same two chemosensory cells pseudocolored blue and yellow. Images show volume reconstructions with (B) or without (C) semi-transparent fill of non-chemosensory cell portions of the cortex, or with only the segmented chemosensory cells shown (D). The volume reconstructions show that each of the chemosensory cells has two microvilli projecting into the environment (green arrows in panel B). See **Supp. Movies 2 & 3** for additional array tomography images of chemosensory cells and their paired microvilli. Scale bar: 2 µm (A).

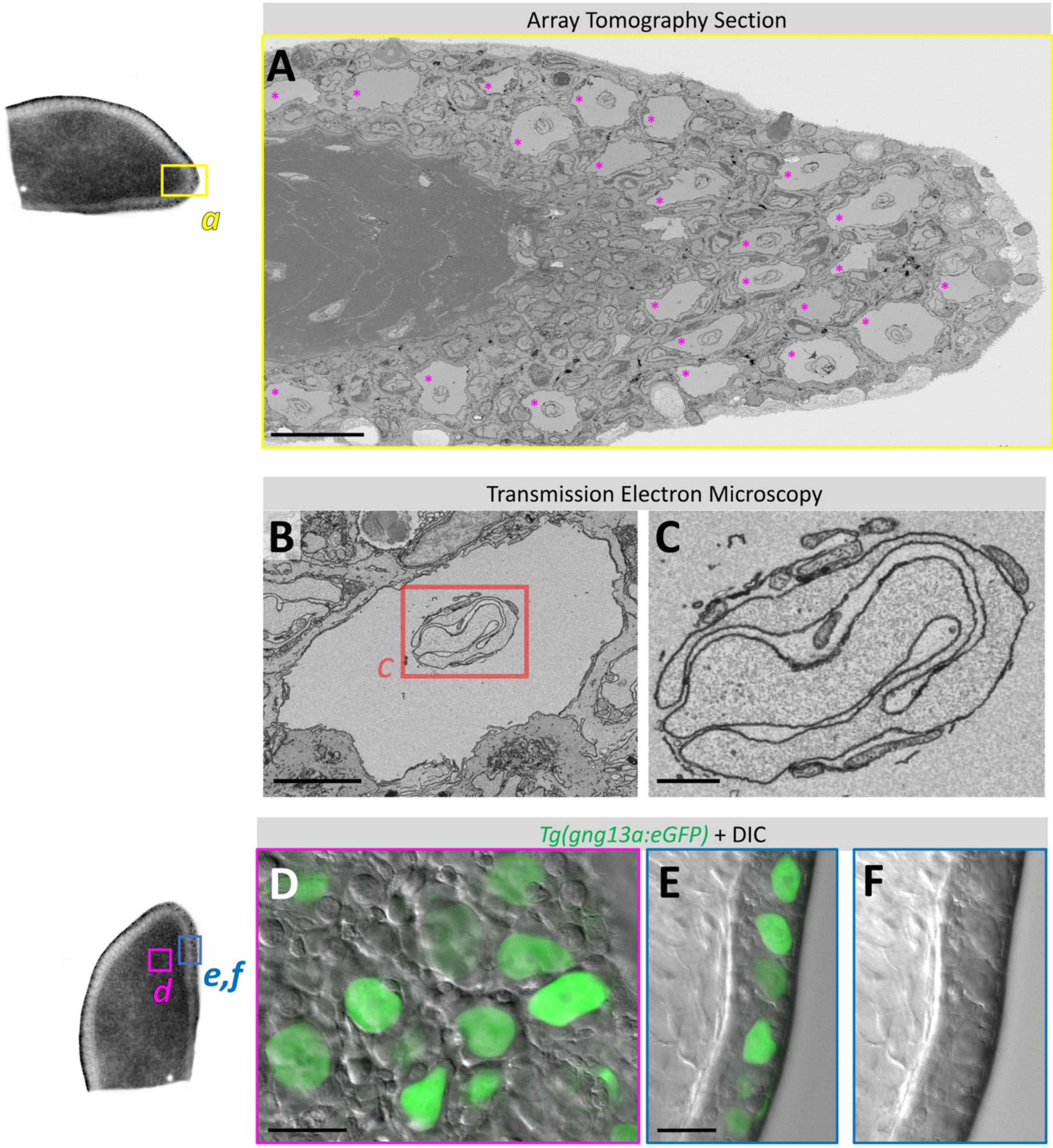


**Fig. S8. Club Cells**

**A.** Single section from an array tomography image stack of an adult zebrafish pectoral axillary ALO, showing an overview tangential section through the edge of a ALO with numerous club cells (magenta asterisks) in the mid-cortex. **B,C.** Transmission electron micrograph of an adult zebrafish pectoral axillary ALO, showing an individual club cell with homogeneous cytoplasm and complex, folded nucleus. Panel C shows a higher magnification image of the nucleus-containing boxed region in panel B. **D-F.** Single section confocal/DIC micrographs of the ALO cortex from a *Tg(gng13a:eGFP)*$^{y709}$ transgenic zebrafish, with *en face* overview (D) and side-view (E) images of the cortex. Panel (F) shows the same image as panel F but with only DIC, showing that the unique club cell morphology, a large cell with a central large nucleus, is easily identifiable through DIC alone. See **Supp. Movie 2** for additional array tomography images of club cells. Scale bars = 20 µm (A), 5 µm (B), 1 µm (C), 25 µm (D,E).

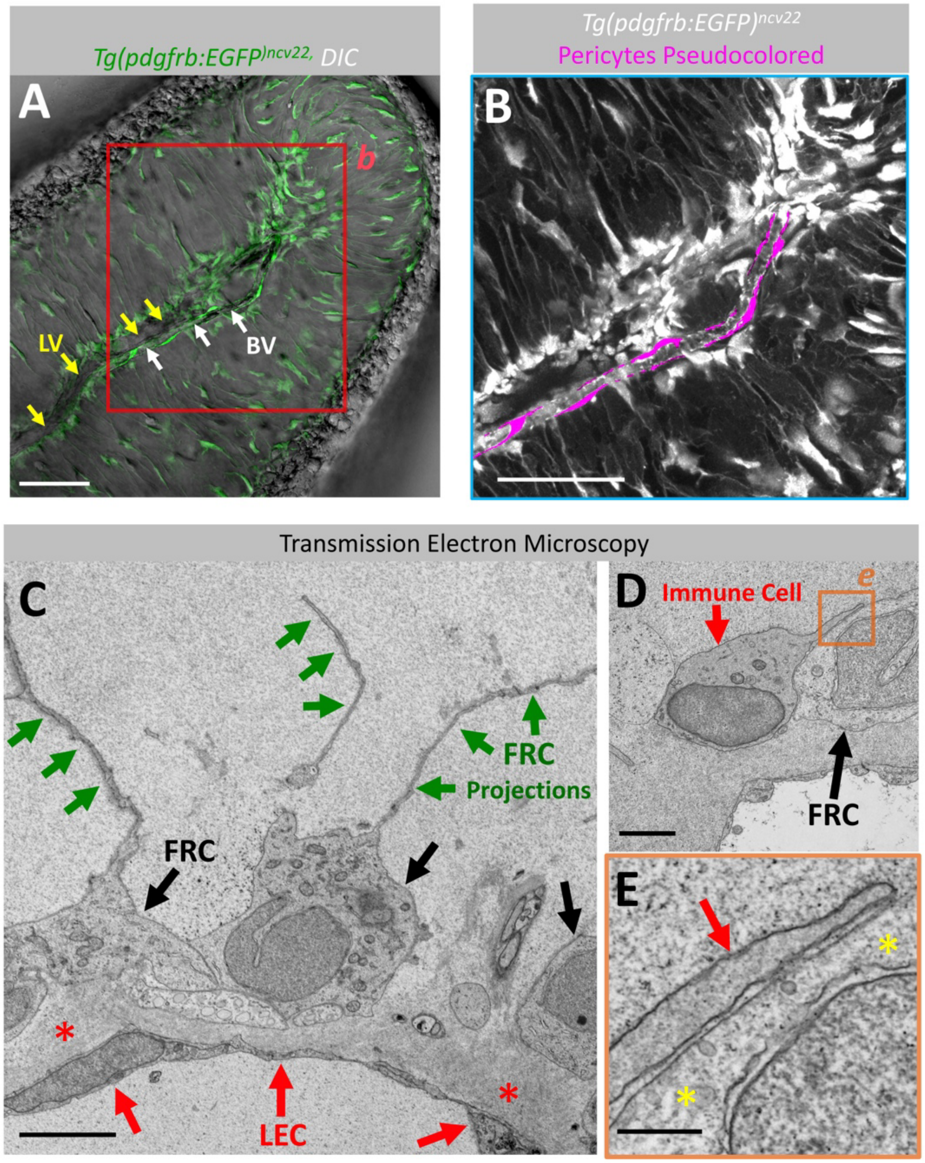


**Fig. S9. Fibroblastic Reticular Cells**

**A-B**. Confocal micrographs of an adult ALO from a *Tg(pdgfrb:EGFP)*$^{ncv22}$ transgenic zebrafish with green fluorescent fibroblastic reticular cells (FRCs) surrounding and extending out from a lymphatic vessel (yellow arrows in panel A), not an adjacent blood vessel (white arrows in panel A). Images show (A) an extended depth of focus image with DIC and (B) a higher magnification image of the boxed region in panel A, showing FRCs in white and pericytes surrounding a blood vessel (also labeled by the transgene) pseudocolored magenta. **C-E.** Transmission electron micrographs of an adult zebrafish pectoral axillary ALO, showing images of FRC's in the ALO. Panel (C) shows three FRC cell bodies (black arrows) surrounding a lymphatic vessel (lymphatic endothelium noted with red arrows), with FRC lamellar extensions noted with green arrows and the thick matrix layer surrounding the vessel noted with red asterisks. Panels D and E show an immune cell (red arrow) closely apposed to a fibroblastic reticular cell body (black arrow) embedded in the matrix surrounding a lympahtic vessel. Panel E shows a magnified image of the boxed region in panel D, with an immune process (red arrow) extending across the FRC (yellow asterisks). See **Supp. Movie 4** for additional images of fibroblastic reticular cells. Scale bars = 50 µm (A,B), 4 µm (C), 2 µm (D), 500 nm (E).

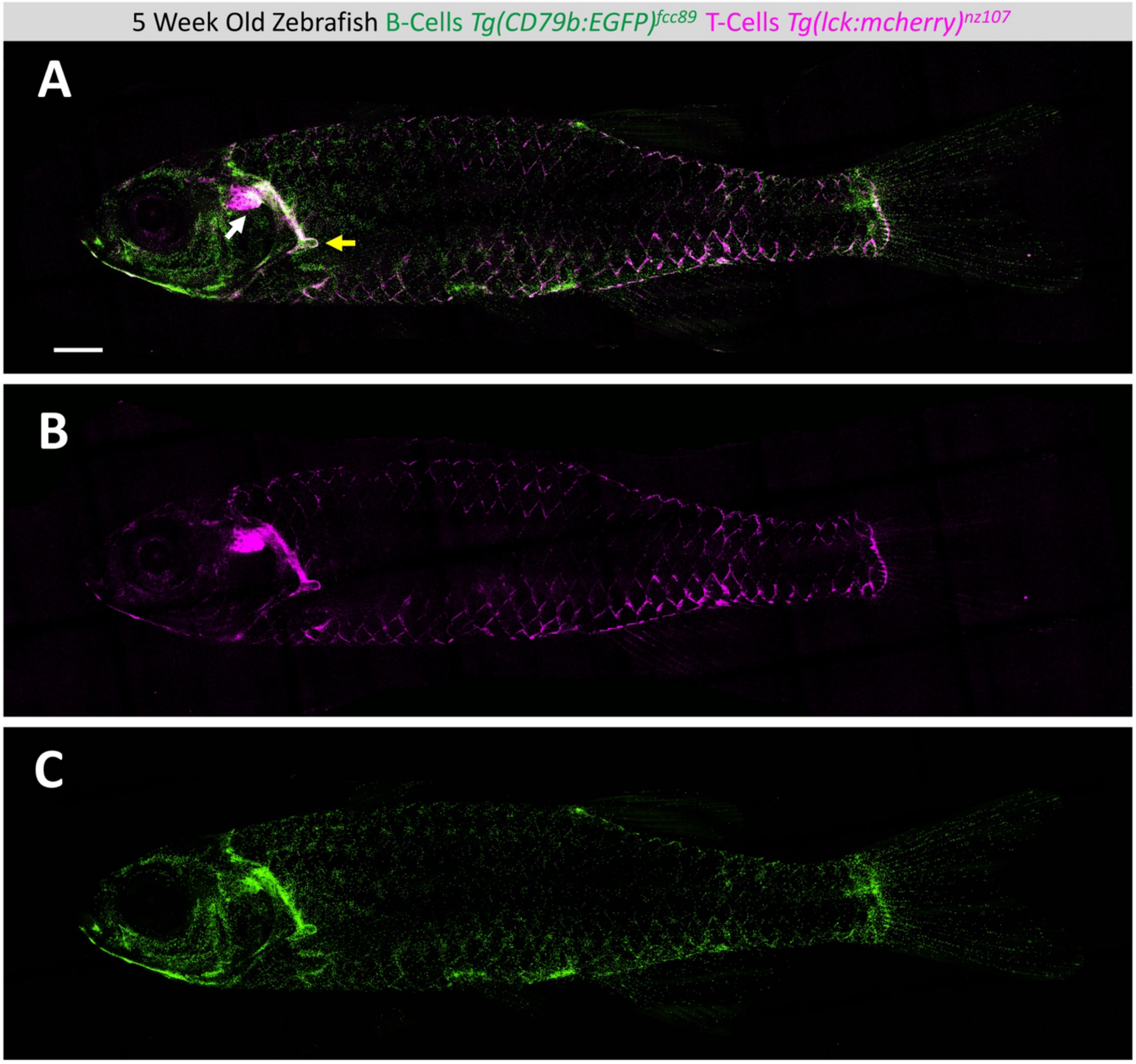


**Fig. S10. The ALO to Thymus Connection**

**A-C.** Confocal micrograph maximum intensity projections of a 5 week-old (16.5 mm standard length) *Tg(cd79b:EGFP)*$^{fcc89}$, *Tg(lck:mcherry)*$^{nz107}$ double transgenic zebrafish, with T-cells in magenta and B-cells in green, showing a connection between the thymus (white arrow) and ALO (yellow arrow). **Fig. 7S-U** shows higher magnification of ALO region. Scale Bar = 1 mm.

**Table S1.** Species of bony fishes (teleosts) examined to assess presence or absence of an ALO-like structure. An ALO like structure is present in the species marked by an asterisk (*). Specimens catalogued in the US National Museum of Natural History are identified by their USNM catalog number. Further information on the USNM specimens is available at https://collections.nmnh.si.edu/search/fishes/

**Teleostei**

Elopomorpha

Elopiformes

Family Elopidae
* *Elops saurus* – USNM 128338

Family Megalopidae
* *Megalops atlanticus* – USNM 303317

Albuliformes

Family Albulidae
*Albula vulpes* – USNM 300474

Anguilliformes
Family Anguillidae
*Anguilla rostrata* – USNM 395727

Notacanthiformes

Family Notacanthidae
*Polyacanthonotus challengeri* – USNM 263242

Family Halosauridae
*Halosaurus ovenii* – USNM 263242

**Osteoglossocephalai**

Osteoglossomorpha

Hiodontiformes

Family Hiodontidae
*Hiodon alosoides* – USNM 350554

Osteoglossiformes

Family Osteoglossidae
* *Osteoglossum bicirrhosum* – USNM 315446

Family Mormyridae
*Marcusenius deboensis* – USNM 357065

Otomorpha

Alepocephali

Alepocephaliformes

Family Alepocephalidae

* *Alepocephalus bairdii* – USNM 215590

Clupei

Clupeiformes

Family Clupeidae
*Clupea harengus* – USNM 325930

Ostariophysi

Gonorynchiformes

Family Chanidae
* *Chanos chanos* - USNM 401640

Family Phractolaemidae
* *Phractolaemus ansorgii* – USNM 203419

Otophysa

Cypriniformes

Cyprinoidei

Family Danionidae
* *Danio rerio* - NIH study material
* *Danio albolineatus* – USNM 390070
* *Danio roseus* – USNM 390510
* *Amblypharyngodon mola* – USNM 344648
* *Rasbora caverii* - USNM 271639
* *Trigonopoma* cf. *agile* – USNM 229233
* *Laubuka* sp. – USNM 271227
* *Esomus altus* – USNM 390719
* *Raiamas nigeriensis* – USNM 339723
* *Nematabramis alestes* – USNM 190116

Family Acheilognathidae
*Acheilognathus lanceolatus* – USNM 50789
*Rhodeus sericeus* – USNM 190163

Family Cyprinidae
*Barbus meridionalis* – USNM 276285
*Carassius auratus* – USNM 191306
*Cyprinus carpio* – USNM 191167
*Tor tambra* – USNM 409895

Family Leptobarbidae
*Leptobarbus hoevenii* – USNM 394013

Family Leuciscidae
*Abramis brama* – USNM 204103
*Agosia chrysogaster* – USNM 377658
*Alburnus alburnus* – USNM 270911
*Rutilus rutilus* – USNM 205936

*Semotiuls atromaculatus* – USNM 382652

Family Tincidae
*Tinca tinca* – USNM 22669

Family Xenocyprididae
* *Chanodichthys erythropterus* – USNM 336724
* *Zacco sieboldii* – USNM 87447

Catostomoidei

Family Catostomidae
*Catostomus macrocheilus* – USNM 369159
*Carpiodes cyprinus* – USNM 374602

Cobitoidei

Family Cobitidae
*Pangio agma* – USNM 328529

Family Balitoridae
*Balitora burmanica* – USNM 378385
*Balitoropsis zollingeri* – USNM 230253
*Hemimyzon taitungensis* – USNM 357418

Family Botiidae
* *Botia histrionica* – USNM 378440

Family Nemacheilidae
* *Paracanthocobitis mandalayensis* – USNM 344646

Family Gastromyzontidae
* *Gastromyzon* sp. – USNM 409794

Family Vaillantellidae
* *Vaillantella euepiptera* – USNM 230272

Gyrinocheiloidei

Family Gyrinocheilidae
*Gyrinocheilus aymonieri* – USNM 117717

Characiformes
Family Characidae
*Astyanax bimaculatus* – USNM 361440

Family Anostomidae
*Leporinus friderici* – USNM 263992

Gymnotiformes
Family Gymnotidae
*Gymnotus carapo* – USNM 225289

Siluriformes
Family Loricariidae
*Hemipsilichthys mutuca* – USNM 342768

Family Sisoridae
*Glyptothorax pictus* – USNM 393760

**Euteleostei**

Lepidogalaxii

Lepidogalaxiiformes
Family Lepidogalaxiidae
*Lepidogalaxias salamandroides* – USNM 401206

Protacanthopterygii

Esociformes
Family Umbridae
*Umbra pygmaea* – USNM 345523

Stomiati
Osmeriformes
Family Osmeridae
*Osmerus dentex* – USNM 401192

Family Retropinnidae
*Retropinna semoni* – USNM 406810

Neoteleostei

Aulopiformes
Family Synodontidae
*Synodus variegatus* – USNM 411546

Percopsiformes
Family Percopsidae
*Percopsis omiscomaycus* – USNM 328315

Gasterosteiformes
Family Gasterosteidae
*Gasterosteus aculeatus* USNM 36982

Beloniformes
Family Adrianichthyidae
*Oryzias latipes* -- USNM 443810

Anabantiformes
Family Badidae
*Badis ruber* – USNM 376436

Tetraodontiformes
Family Tetraodontidae
*Takifugu rubripes* -- USNM 49839

**Movie S1. ALO regeneration**
A combination of still images and live video showing ALO regeneration over fourteen days, taken with a stereomicroscope.

**Movie S2. Array tomography overview and ALO cortex scan**
Array tomography image Z stack through the ALO, with a higher magnification scan of the ALO cortex.

**Movie S3. Array tomography of chemosensory cells and extruding goblet cells**
Array tomography image Z stack highlighting chemosensory cells with two microvilli extending into the environment, as well as goblet cells extruding mucus, followed by 3D reconstructions with chemosensory cells segmented.

**Movie S4. Adult ALO vasculature**
3D reconstruction and real-time single-plane live video of confocal micrographs of ALOs on adult *Tg*(*mrc1a:egfp)*$^{y251}$, *Tg*(*kdrl:mcherry)*$^{y205}$ double transgenic zebrafish injected intravascularly with Hoechst 33342 dye, with lymphatic vessels in green, blood vessels in magenta, and circulating Hoechst dye-labeled red blood cell nuclei in blue.

**Movie S5. Fibroblastic reticular cells**
Scroll through Z and 3D reconstructions of ex-vivo confocal + DIC imaged ALO from an adult transgenic *Tg(pdgfrb:egfp)*$^{ncv22}$ transgenic zebrafish, with fibroblastic reticular cells in green, followed by an array tomography scroll through Z highlighting an immune cell and a fibroblastic reticular cell.

**Movie S6. Immune cells migrate on Fibroblastic Reticular Cells**
*Ex vivo* confocal timelapse extended depth of focus imaging of a ALO from an adult *Tg(mrc1a:egfp)*$^{y251}$ transgenic zebrafish, with a few macrophages in green as well as many other immune cells migrating along the fibroblastic reticular cell network.

**Movie S7. Live imaging of macrophages in the adult zebrafish ALO**
*Ex vivo* confocal micrograph extended depth of focus and maximum intensity projection timelapse movies of the ALO of an adult *Tg(mpeg1:egfp)*$^{gl22}$ transgenic zebrafish with macrophages in green.

**Movie S8. Live imaging of B cells in the adult zebrafish ALO**
*Ex vivo* confocal micrograph extended depth of focus and maximum intensity projection timelapse movies of the ALO of an adult *Tg(cd79b:egfp)*$^{fcc89}$ transgenic zebrafish with B-cells labeled in green.

**Movie S9. Live imaging of T cells in the adult zebrafish ALO**
Ex vivo Confocal micrograph extended depth of focus and maximum intensity projection timelapse movies of the ALO of an adult *Tg(lck:egfp)*$^{cz2}$ transgenic zebrafish with T-cells labeled in green.

**Movie S10. The ALO in T-cell acute lymphoblastic leukemia**

*Ex vivo* confocal micrograph scroll through Z of ALOs of adult *Tg(lck:egfp)*$^{cz2}$ transgenic wild-type sibling or HLK$^{dz102}$ transgenic fish with T-cell acute lymphoblastic leukemia. Normal or leukemic T-cells are labeled in green by *lck:egfp* expression.
